# A developmental gradient reveals biosynthetic pathways to eukaryotic toxins in monocot geophytes

**DOI:** 10.1101/2023.05.12.540595

**Authors:** Niraj Mehta, Yifan Meng, Richard Zare, Rina Kamenetsky-Goldstein, Elizabeth Sattely

## Abstract

Numerous eukaryotic toxins that accumulate in geophytic plants are valuable in the clinic, yet their biosynthetic pathways have remained elusive. A lead example is the >150 Amaryllidaceae alkaloids (AmAs) including galantamine, an FDA-approved treatment for Alzheimer’s disease. We show that while AmAs accumulate to high levels in many tissues in daffodils, biosynthesis is localized to nascent, growing tissue at the base of leaves. A similar trend is found for the production of steroidal alkaloids (e.g. cyclopamine) in corn lily. This model of active biosynthesis enabled elucidation of a complete set of biosynthetic genes for the production of AmAs. Taken together, our work sheds light on the developmental and enzymatic logic of diverse alkaloid biosynthesis in daffodil. More broadly, it suggests a paradigm for biosynthesis regulation in monocot geophytes where plants are protected from herbivory through active charging of newly formed cells with eukaryotic toxins that persist as aboveground tissue develops.

## Introduction

Geophytes are a large and diverse group of plant life-forms with underground food-storage organs that comprise various species of important edible, ornamental and medicinal crops^1^. Their cycles of dormancy and active growth are governed by environmental conditions, e.g., temperature and photoperiod. Under unfavorable conditions, plants enter dormancy, bulbs remain underground without leaves and roots, and young leaves and flowers develop inside the bubs at the expense of storage tissue. Frequently, therefore, geophytes produce a battery of chemical toxins and repellants to defend these food reserves from herbivores. For example, in addition to their culinary uses, organosulfur flavor-compounds produced by the bulbous geophytes onion and garlic are potently repellant to insects and animals^2, 3^. Geophytes’ potent bioactivities towards herbivores and other eukaryotes have served as a reservoir of medicinal metabolites that can modulate human biology as well – for example, the steroidal glycoalkaloids (e.g. cyclopamine) produced by the rhizomatous, bulbous Veratrum genus of plants inhibit the Hedgehog signaling pathway, and thus find applications in treating various cancers^2–6^. Similarly, *Convallaria* and *Urginea* contain potent cardioactive glycosides that are toxic to herbivores^7–12^. Although these geophyte toxins are a rich source of bioactive metabolites for health and agronomic applications, very few of the biosynthetic pathways that are involved in the production of these toxins have been described^13, 14^.

Few geophytes produce as remarkable and extensive a range of bioactive metabolites as do plants in the Amaryllidoideae subfamily of monocots, which includes common household plants such as daffodils^15, 16^. Collectively known as the *Amaryllidaceae* alkaloids (AmAs), these metabolites exhibit extensive structural and functional diversity (Figure 1)^17–20^. AmAs include the FDA-approved drug galantamine, which is of the benzofurobenzazepine class. Galantamine, isolated from *Narcissus* (daffodil), *Lycoris* (red spider lily), and other Amaryllidoideae species, is a cholinergic modulator that is one of only five therapeutics approved for use in the treatment of cognitive symptoms in Alzheimer’s Disease (AD)^21^. In daffodils, many other structurally diverse, but biogenically related alkaloids co-occur with galantamine. Haemanthamine, for example, is an AmA built from the same precursors as galantamine in daffodil bulbs but possesses a 5,10b-ethanophenanthridine skeleton. Haemanthamine exhibits no cholinergic modulation and instead inhibits eukaryotic cell growth by direct binding to the ribosome^22, 23^. Narciclasine, another alkaloid from daffodils, possess an isocarbostyril skeleton and exhibits potent, nanomolar-range eukaryotic growth inhibition through inhibition of protein synthesis, in addition to modulating, through direct binding, the eukaryotic translation elongation factor eEF1A^24–26^. Lycorine, one of the most abundant AmAs in daffodils, is of the pyrrolophenanthridine type and exhibits eukaryotic growth inhibition. In addition, lycorine displays significant antiviral activity against flaviviruses such as West Nile viruses, the dengue virus and has been reported to inhibit SARS-CoV-2 replication through inhibition of its Main protease (M^pro^)^27–31^. The biosynthetic pathways to AmAs are largely unknown and, to date, only three enzymes have been associated with the pathway^32–34^ Although the over 150 known AmAs possess extensive structural and functional diversity, they are all produced through a relatively small set of biochemical transformations of a common amino-acid derived precursor, 4-O’-methylnorbelladine, 4OMN. As such, this class of alkaloids represents a striking natural example of *diversity oriented biosynthesis, i.e.* the parsimonious generation of chemical diversity through iterative diversification of the same precursors. Understanding how this diversification occurs would shed insight on how nature achieves such efficient chemical and functional diversification of simple precursors. Furthermore, discovering the genetic basis for AmA biosynthesis would permit a finer understanding of their native biological roles and open doors to their sustainable production in scalable heterologous systems like yeast and *Escherichia coli*, thus bypassing supply challenges that limit their clinical development^35, 36^.

**Figure 1:**
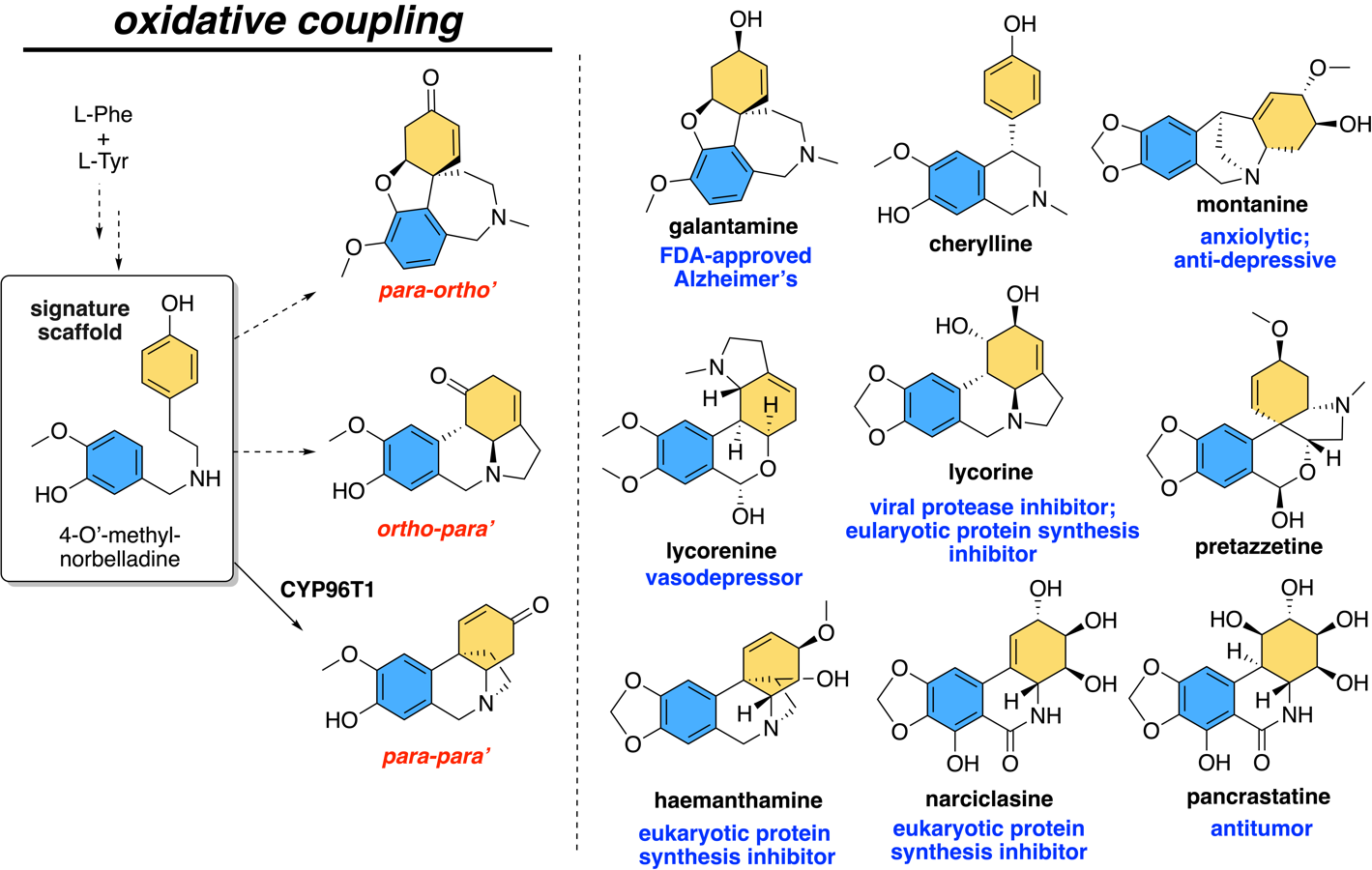
The Amaryllidaceae Alkaloids (AmAs) are a biogenically related set of structurally diverse bioactive alkaloids. 4-O’-methylnorbelladine (4OMN), a 2° amine derived from L-Phe and L-Tyr, serves as the precursor to a large diversity of bioactive alkaloids. Reported bioactivities are listed in blue font^22, 23, 27, 102, 120–133^.

Prior labelling studies have shed light on the chemical logic of AmA biosynthesis and diversification^17–19, 37–43^. Early steps of the biosynthesis of the Amaryllidaceae alkaloids involve the production of the dialkalyamine norbelladine through reductive condensation of the L-Phe derived aldehyde 3,4-dihydroxybenzaldehyde with the L-Tyr derived amine, tyramine^18, 41^. Subsequent methylation at the 4’-hydroxyl of norbelladine affords 4-O’-methylnorbelladine (4OMN), which serves as the immediate precursor to the subsequent C-C oxidative coupling step that constitutes the first committed step in AmA biosynthesis (Supplementary Figure S1)^18, 38, 39^. Differential oxidative coupling of this linear precursor gives rise to the three basic scaffold types that comprise the vast majority of Amaryllidaceae alkaloids^44^. These three basic scaffold types, named after the regiochemistry of oxidative coupling that affords their scaffolds, i.e. *para-para’* (*p-p’*, e.g. haemanthamine), *ortho-para’* (*o-p’,* e.g. lycorine), and *para-ortho’* (*p-o’,* e.g. galantamine), are then diversified into an elaborate set of AmAs through subsequent oxidations, methylations and ring-rearrangements^16^. As such, identifying enzymes for this gatekeeping oxidative coupling of 4OMN is key to understanding the genesis of structural diversity in the AmAs. Previous work has revealed one cytochrome P450 (P450) enzyme, CYP96T1, that affords the *para-para’* scaffold, but additional enzymes for the remaining two modes of oxidative coupling have not been described^33^.

Many plant pathway discovery efforts lean heavily on transcript co-expression to identify candidate biosynthetic enzymes, and there have been several efforts for large-scale sequencing of medicinal plants of interest^45, 46^. In the case of the AmAs, multiple producing plants have been analyzed^28, 42, 43^. However, in these datasets, CYP96T1 was not found to be highly expressed in the sequenced tissues, making identification of other enzymes difficult. (Supplementary Figure S2, Supplementary Figure S3) It is also known that the AmAs are distributed in high concentrations in many parts of the plant, potentially obscuring their site of origin^49–51^. Here, we use a combination of precursor feeding studies, metabolomics, and transcriptomics of carefully sectioned tissue to determine a site of AmA biosynthesis in daffodil tissues. Long read RNA-seq methods were essential for these polyploid species in order to deconvolute similar transcripts that encode enzymes with different activity. This effort enabled the identification of a series of candidate genes that could be tested through heterologous expression in *Nicotiana benthamiana* using Agrobacterium-mediated transient T-DNA expression for activity in the AmA pathway. Ultimately this approach unveiled an elegant strategy to metabolic diversification in a class of ornamental plants with direct relevance to human health.

## Results

### Labelling studies identify leaf bases as the sites of active biosynthesis

Broadly, the AmA biosynthetic pathway consists of three major branches— the *ortho-para’* (e.g. lycorine), *para-para’* (e.g. haemanthamine) and *para-ortho’* (e.g. galantamine) pathways, each named after the regiochemistry of oxidative coupling occurring at the first step of these pathways (Figure 1). While several transcriptomic and metabolite analyses have previously been reported for various Amaryllidoideae species, enzymes for only the *para-para’* pathway have been identified. These enzymes include a key oxidative coupling enzyme CYP96T1, which generates the *p-p’* scaffold, and a reductase that generates a metabolite that is side product in haemanthamine biosynthesis^33, 34^. It is unknown whether the other two pathways, i.e. *p-o*’ and *o-p’*, are also co-expressed and co-regulated with the *para-para’* pathway inside the bulb. For non-model plant like daffodils, where genomic information and genetic tools are often unavailable, RNA-seq expression profiling of biosynthetically active tissue is the principal approach to identifying candidate pathway enzymes^52–54^. However, sequencing a plant transcriptome without knowing where a pathway is expressed may result in significant under-sampling of pathway genes^55^. Therefore, we first sought to understand the patterns of alkaloid accumulation and production across each of the three distinct biosynthetic pathways.

In initial efforts to study metabolite accumulation, we first compared abundance of key alkaloids from each of the three AmA pathways (*p-p’*, *p-o’*, *o-p’*) in different tissues of daffodils *Narcissus* cv. Tête-à-Tête (*Narcissus cyclamineus × Narcissus ’Cyclataz’*) at full bloom using liquid chromatography mass spectrometry (LC-MS). Key metabolites from each of the three pathways were detected at high levels in several tissues, but in some cases, their accumulations differed from each other (Supplementary Figures S4 and S5), consistent with previous reports in other *Narcissus* species^49, 50^. Either these metabolites 1) are being actively made in the tissues that they were being detected in, or 2) were distributivity transported to their sites of accumulation from a different source tissue where they were being actively produced.

In an effort to differentiate between these two possibilities and capture the site of active biosynthesis in an RNA-seq experiment, we next directly measured *active* biosynthesis in different parts of the plant. In daffodil leaves, cell division and elongation occurs within the bulb, therefore, the youngest sections of the leaf are located at the leaf bases present in the core of the bulb organ^56^. We sectioned different daffodil tissues, including bulb scales and foliage tissue, and soaked them in a solution of isotopically labelled precursor, [^2^H]-4-O’-methylnorbelladine ([^2^H]-4OMN). The labelled di-alkylamine [^2^H]-4OMN is the common substrate for the first committed steps for each of the three AmA pathways. Its incorporation into downstream AmA metabolites would therefore allow us to identify whether each of the three biosynthetic pathways were active in a given tissue. After treatment, we analyzed metabolite extracts by Liquid Chromatography Mass Spectrometry (LC-MS) in an untargeted fashion for labelled downstream AmA pathway metabolites that were significantly enriched in abundance in tissues fed with the substrate relative to a water control.

Daffodil leaf sections natively contained high levels of key alkaloids across all three pathway branches, namely haemanthamine (*p-p’*), lycoramine (*p-o’*) (i.e. galantamine with a reduced alkene) and lycorine (*o-p’*). The highest levels of these natively accumulated alkaloids were in the oldest parts of the leaves, which correspond to the very top of the leaves (Figure 2a). Conversely, the lowest levels of these alkaloids were found in the youngest, newly developed parts of the leaves, closest to their bases that were underground within the bulb. Remarkably, however, when we tracked active biosynthesis through isotope labelling, we observed a complete inversion of this trend— active biosynthesis, as measured by tracking production of the labelled *p-p’* pathway intermediate [^2^H]-vittatine (i.e. a haemanthamine precursor), was highest in the bottommost (i.e. youngest, actively growing) parts of leaves and decreased progressively towards the topmost (i.e. oldest) parts of the leaves. Thus, although the youngest sections of the leaves accumulate the lowest levels of accumulated alkaloid toxins, they exhibit the highest rates of their active biosynthesis (Figure 2). We also showed using reverse transcription polymerase chain reaction (RT-PCR), relative to a histone control as internal reference, that the expression of the known pathway gene CYP96T1 matched the isotope labelling pattern. These data suggest that isotope labeling patterns are indeed indicative of active pathway metabolism. Moreover, this trend held true for all three branches of the pathway, *p-p’*, *p-o’*, and *o-p’*, indicating that all AmA biosynthesis is co-localized within the same tissue (Figure 2b). 2-Dimenional Desorption Electrospray Ionization Mass Spectrometry (DESI-MS) imaging data on similarly fed tissues in paperwhite daffodils (*Narcissus papyraceus)* further corroborated our findings, revealing that more labelled intermediate can be detected in intact leaf base tissue than in older tissue (relative to unlabeled lycorine). Interestingly, the images revealed that AmAs may be localized proximal to vascular tissue (see distributed puncta in transverse cross-sections of daffodil leaves in Figure 2c, Supplementary Figures S6, S7 and S8).

**Figure 2:**
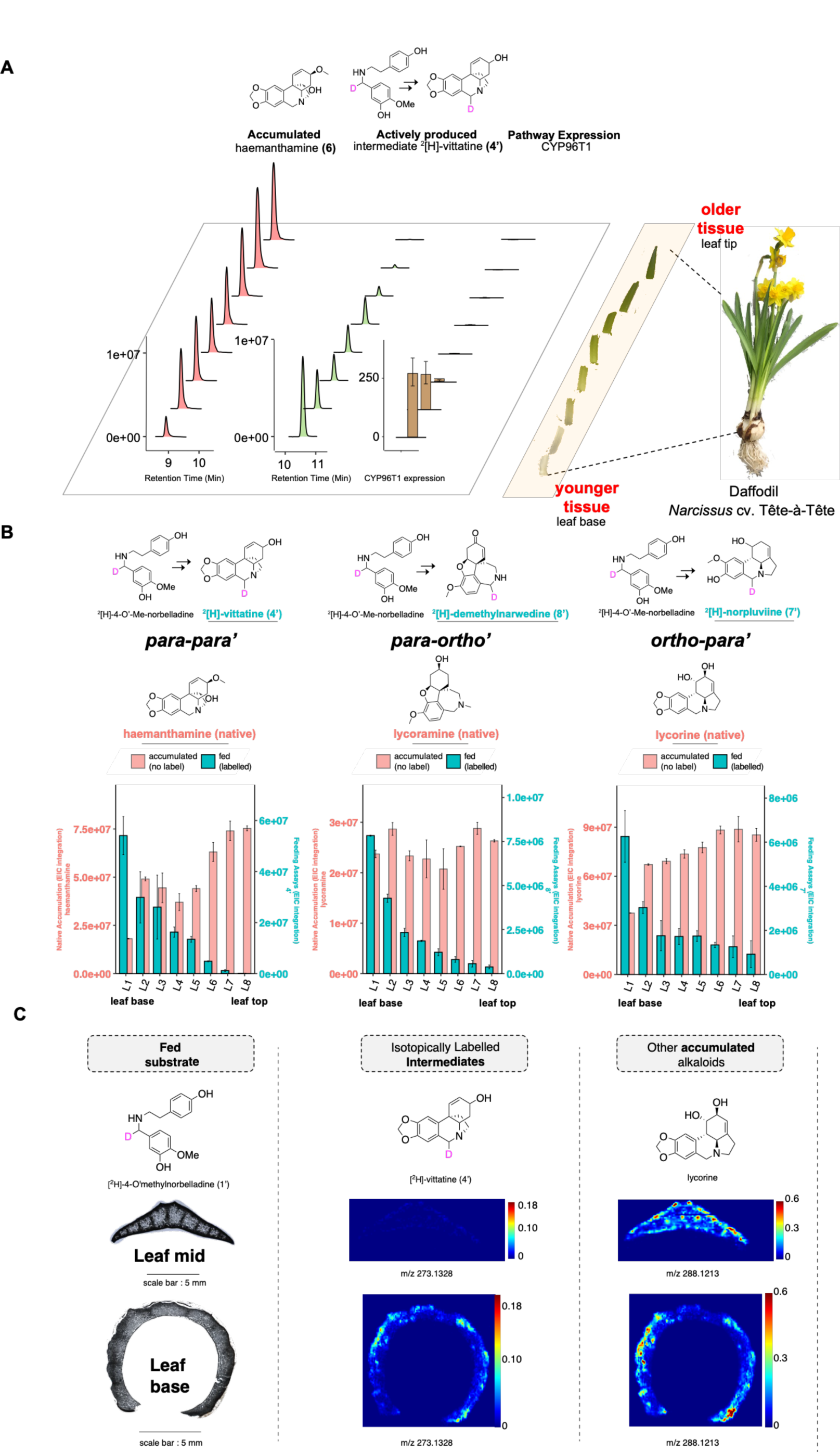
Stable-isotope precursor feeding experiments identify leaf bases as the sites of active biosynthesis. **A)** Leaf tissues from a fully-grown daffodil plant at a peak blooming stage were sectioned for assessing levels of the highly-accumulating alkaloid haemanthamine through LC-MS. In a parallel experiment from the same plant, similarly-sectioned leaf tissues were individually fed with the haemanthamine precursor [^2^H]-4-O’-methylnorbelladine, and incorporation into the haemanthamine pathway intermediate vittatine (Supplementary Figure S19) was tracked to measure active biosynthesis. The expression of the haemanthamine pathway gene *Nt*CYP96T1 was also quantified in parallel on freshly-harvested leaf sections from the same plant. Daffodils are monocotyledonous plants that exhibit bottom-up leaf tissue growth, such that the bottom-most sections of the foliage leaves are the youngest parts of the leaves. While the levels of un-labelled, natively accumulated haemanthamine are highest in fully mature, older leaf tissue sections and lowest in nascent, young basal leaf tissue, active biosynthesis of those metabolites is highest in the young, basal leaf tissue, and lowest in older leaf tissue. **B)** All three branches of the AmA pathways exhibit the same trend observed in (A) – active biosynthesis is highest in the basal leaf tissues (section L1, Supplementary Figure S29) even though accumulation is highest in the older, mature leaf tissues (e.g. section L8). The extracted ion abundances (peak integration) are shown for: unlabeled haemanthamine ([(M+1)+H]^+^ = *m/z* 303.1426), unlabelled lycoramine ([(M+1)+H]^+^ = *m/z* 291.1790), unlabelled lycorine ([(M+1)+H]^+^ = *m/z* 289.1270 [(M+1)+H]^+^), deuterium labelled ^2^[H]-norpluviine ([M+H]^+^ = *m/z* 275.1506), deuterium labelled ^2^[H]-demethylnarwedine ([M+H]^+^ = *m/z* 273.1349), deuterium labelled ^2^[H]-vittatine ([M+H]^+^ = *m/z* 273.1349). N=2 biological replicates for all feeding and metabolite quantification experiments, and N=1 biological replicates (3 technical replicates) for quantifying *Nt*CYP96T1 expression levels as previously reported^47^, using the Histone gene for internal reference. The expression fold change was quantified using the 2^-ΔΔCt^ method. All experiments measuring the accumulation of un-labelled intermediates here were run on separate sets of leaves from the same plant and were not fed any precursors. The M+1 isotopologue for the natively accumulated, unlabelled alkaloids haemanthamine, lycoramine and lycorine were quantified to avoid detector saturation effects, as these alkaloids were present in very high abundances in our measurement conditions. **C)** DESI-MS images for transverse leaf sections of *Narcissus papyraceus* (paperwhites). Intact leaf tissue that was split into two halves via a transverse cut (top half and bottom half) and each half was fed deuterium labelled precursor ([^2^H]-4OMN) for 7-14 days. The basal ∼1 inch of tissue was then sectioned on a cryostat. The mass ion intensities for each sample shown here were normalized to the maximum ion intensity of lycorine within the same sample. Scale bars shown.

The annual cycle of Narcissus consists of stages of dormancy and active growth. In cooler regions, as temperatures rise in spring, the bulb breaks out of dormancy and new leaves start emerging from within the bulb. Leaf elongation begins with cell division at the bases of these newly emerging leaves located inside the Narcissus bulb^56, 57^. Collectively, our feeding data suggest a model where these young, actively growing leaves are charged with alkaloid toxins before they emerge above-ground for photosynthesis, thus revealing a logic for toxin biosynthesis in the Amaryllidoideae and possibly other geophytes. Furthermore, these data indicate that capturing this early developmental stage under active growth conditions is critical for finding pathway enzymes. With the identification of young leaf bases as the sites of active biosynthesis, we next turned to candidate gene identification through RNA-seq expression profiling of biosynthetically active tissue.

### Long-read transcriptome Sequencing enables identification of candidate P450s for oxidative coupling

Having identified leaf bases as sites of active biosynthesis, we performed RNA-seq on various tissue segments of daffodils, including young leaf bases and older leaf tips, to allow for identification of candidate pathway genes using expression analysis (Supplementary Table S1). We paired our RNA-seq experiment with isotope labelling studies on leaves from the same plant to ensure that we had correctly chosen biosynthetically active tissue for our experiment. We complemented our dataset with long-read Pacbio IsoSeq to overcome difficulties in assembling the polyploid (allotriploid) transcriptome of Tête-à-Tête daffodils from short-read RNAseq data alone and to generate an accurate, full-length transcriptome assembly^58^. Encouragingly, we observed that the previously characterized pathway enzyme CYP96T1 was amongst the most highly expressed cytochromes P450 transcripts in the leaf base tissues, suggesting that our transcriptome sequencing had captured the relevant biosynthetic tissue (Supplementary Figure S2). This highlights the importance of direct sampling of biosynthetically active tissue for RNA-seq experiments – the expression of CYP96T1 was found to be much lower in other, publicly reported transcriptomes that sampled from much larger sections of bulb tissue, including presumably, parts not active in AmA biosynthesis (Supplementary Figure S2).

With a quality transcriptome assembly (BUSCO 89.1% completeness) that had captured the sites of active biosynthesis, we first sought to identify enzymes for the gatekeeping steps of AmA biosynthesis, i.e. the oxidation of 4OMN in one of three different ways to generate the *para-ortho’* (*p-o*’), *ortho-para’* (*o-p’*), and *para-para*’ (*p-p’*) scaffolds, from which all subsequent chemical diversity results (Figure 3A).

**Figure 3:**
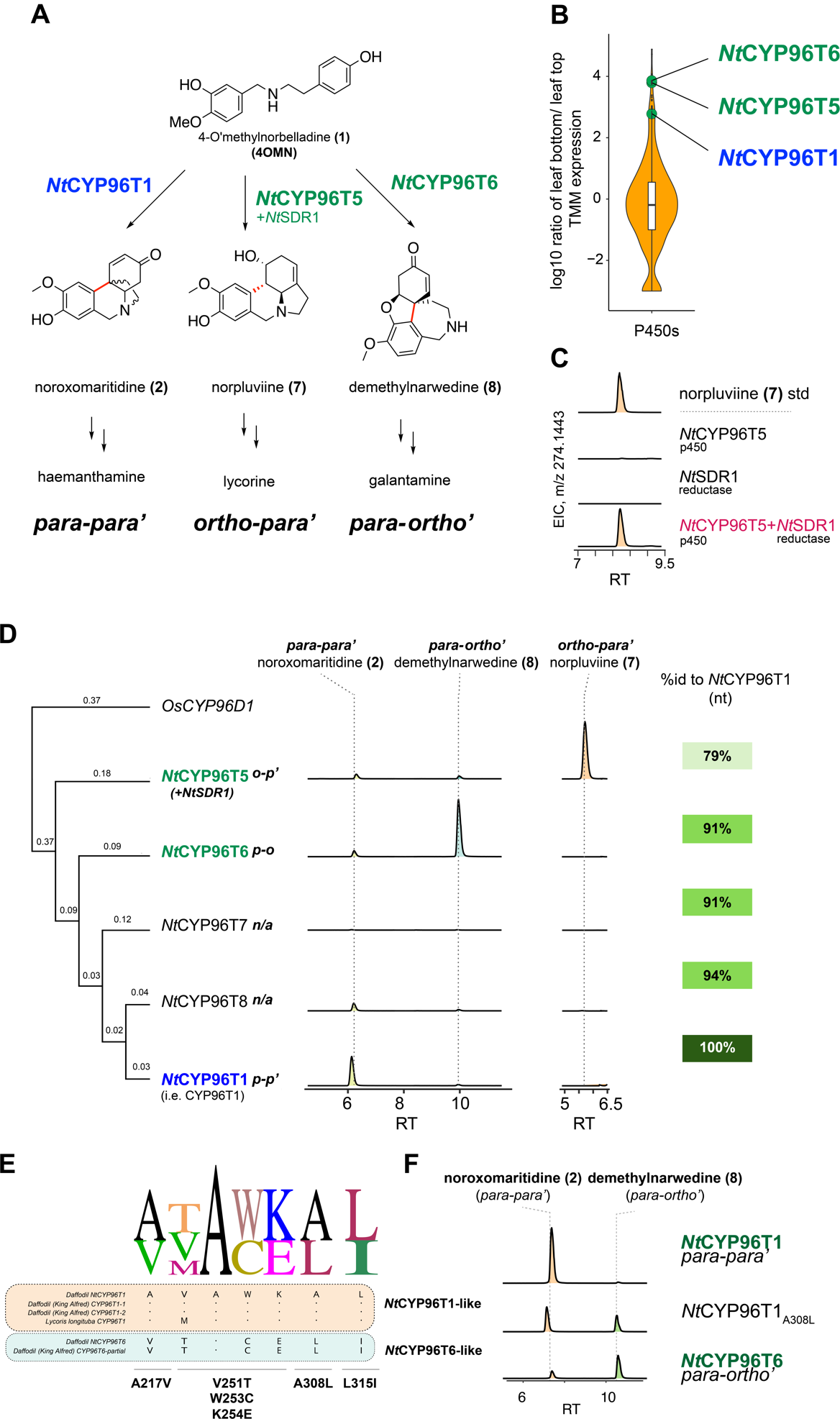
Three highly similar oxidative-coupling Cytochrome P450 homologs are key to diversity generation in the AmAs. **A)** The differential oxidative coupling of 4-O-methylnorbelladine into products with the *para-ortho’* (*p-o*’), *ortho-para’* (*o-p’*), and *para-para*’ (*p-p’*) regiochemistries constitutes the first committed step of AmA biosynthesis (panel A). Whilst the enzyme for the *p-p’* mode of oxidative coupling to noroxomaritidine has been previously reported (CYP96T1), the enzymes for the other two modes of coupling remained unknown. **B)** We searched for candidate oxidative coupling enzymes by looking for leaf-base expressed Cytochromes P450 that exhibit high expression in leaf bases and low expression in leaf tips. By testing candidate genes through *Agrobacterium*-mediated transient T-DNA expression in *N. benthamiana*, we identified a 91% CYP96T1 homolog (nucleotide level), *Nt*CYP96T6 that can catalyze the *p-o’* oxidative coupling and another 79% CYP96T1 homolog (nucleotide level), *Nt*CYP96T5 that can catalyze the *o-p’* coupling**. C)** While *Nt*CYP96T5 alone does not produce the expected oxidative coupling product noroxopluviine, *Nt*CYP96T5 and the short-chain dehydrogenase/reductase *Nt*SDR1, when expressed with *Nt*CYP96T5, produces the *ortho-para’* pathway intermediate norpluviine. **D)** A phylogenetic tree of representative CYP96T1-like sequences in our *Narcissus* cv. Tête-à-Tête transcriptome, and extracted ion chromatograms for the oxidative coupling activities for each of the enzymes. *Nt*CYP96T6 (*p-o’*), *Nt*CYP96T5 (*o-p’*) and *Nt*CYP96T1 (*p-p’*) have different regioselectivites for each of the three branches of the pathway. *Os*CYP96D1: *Oryza sativa* CYP96-family protein was used as an outgroup. **E)** We sought to identify residues that imparted *Nt*CYP96T1 and *Nt*CYP96T6 differential *p-p’* and *p-o’* regioselectivity by looking for conserved residue differences within the Substrate Recognition Sites of these P450s across their homologs in other AmA producer species. We identified a set of four candidate regioselectivity-determining sites on these P450s. **F)** Inversion site-directed mutagenesis of a candidate regioselectivity-determining site (from panel (E)) reveals that swapping Alanine 308 on the *p-p’* coupling enzyme CYP96T1 to the corresponding Leucine on the *p-o’* coupling enzyme *Nt*CYP96T6 imparts *Nt*CYP96T1 the ability to catalyze *p-o’* oxidative coupling like *Nt*CYP96T6.

In searching for enzymes that could oxidatively couple 4-O’-methylnorbelladine (4OMN) to the *p-o’* and *o-p’* intermediates demethylnarwedine (**8**) and noroxopluviine respectively, we focused our efforts on identifying cytochromes P450 (i.e. P450) candidates, which are a family of plant enzymes known to catalyze a broad range of oxidative reactions, including C-C oxidative coupling^33, 59^. A total of 33 unique P450s with high expression in leaf bases and low expression in leaf tops were selected for study (Figure 3B, Supplementary Figure S2). Curiously, we noted that the list was heavily dominated by transcripts of the CYP96-family; several had particularly high homology to CYP96T1 (>80% nucleotide identity) (Supplementary Figure S9). Since CYP96T1 has already been shown to accept 4OMN as a substrate to produce the *p-p’* type product noroxomaritidine (**2**), we reasoned that these CYP96T1-homologs might have evolved to catalyze alternate modes of oxidative coupling from a common substrate. Indeed, small changes to an enzyme’s amino-acid composition can significantly affect its catalytic regioselectivity^60–62^. For example, the *Arabidopsis thaliana* P450s CYP71A12 and CYP71A13, while 89% identical to each other at the amino acid level, oxidize the same substrate to produce different products in camalexin biosynthesis^63^.

We therefore cloned and tested each of these P450s, and all other CYP96T1 homologs within the transcriptome, using *Agrobacterium*-mediated transient T-DNA expression in *Nicotiana benthamiana* leaves and direct co-infiltration of the 4OMN substrate (Methods, Supplementary Figures S9 and S10)^64^. Analysis of infiltrated leaf extracts by LC/MS confirmed that, as expected, CYP96T1 produced one major peak with the expected MS/MS fragmentation pattern for **2**, the *p-p’* product (Figure 3D)^33^. Most of the other CYP96T1-like candidates either produced the same *p-p’* product, or alternatively, no oxidized products with the expected [M+H]^+^ = *m/z* 272.1281 could be identified (Figure 3D). However, one CYP96T1-like P450, *Nt*CYP96T6, a homolog of CYP96T1 with 91% identity at the nucleotide level, produced the galantamine-type *p-o’* intermediate **8**,^65^ which upon methylation and reduction of the carbonyl, would afford galantamine.

### Identification of Oxidative Enzymes for the *ortho-para’* mode of oxidative coupling

Since both enzymes for the oxidative couplings of 4OMN to the *p-p’* and *p-o’* products were closely related P450s of the CYP96 family, we speculated that the enzyme for the third, i.e. *o-p’* mode of oxidative coupling would also be a closely related P450 of the same CYP96 family. However, despite extensive testing of candidate P450s, we were not able to identify this enzyme after expressing candidates in the heterologous *N. benthamiana* testing system. This was puzzling— the *ortho-para*’ AmA lycorine is one of the most abundant alkaloids in daffodils,^49, 51^ and our feeding studies had suggested that it was being actively made in leaf bases (Figure 3B), where we had determined that the *p-p’* and *o-p’* pathways are active.

Several explanations could explain our inability to identify the requisite enzymes. These range from poor chemical stability of enzyme products to poor protein expression of the requisite enzyme in the heterologous host *N. benthamiana*. Without a clear hypothesis for the candidate enzyme type, we speculated that since we observed active biosynthesis in our precursor feeding experiments in daffodil leaf bases, the relevant set of enzymes for lycorine biosynthesis should also be present in the same list of leaf-base expressed enzymes in which we had found biosynthetic genes for the *p-p’* and *p-o’* pathways. We considered that if all enzyme candidates were transiently expressed into the same *N. benthamiana* leaf at once, we might be able to identify the right set of enzymes to at least partly reconstitute the lycorine pathway in *N. benthamiana* leaves.^66, 67^

We identified, cloned and simultaneously co-infiltrated Agrobacterium strains carrying T-DNA for enzyme-encoding genes that are highly co-expressed with *Nt*CYP96T1. These candidates were tested in *N. benthamiana* leaves in different combinations grouped by putative enzyme function; infiltrations contained up to 26 candidates at a time in the same leaf (Methods, Supplemental Figure S11). Remarkably, even when 25 candidates were co-infiltrated, we were able to clearly detect production of new metabolites (Supplemental Figure S12). Since we were infiltrating multiple types of enzymes in the same leaf, we did not know what pathway intermediates would be produced, if any, and so we conducted a semi-targeted search, looking for mass features that corresponded to the expected *m/z* values for putative pathway intermediates identified in prior labelling studies^19, 37, 38, 42, 43^. In one set of batch co-infiltration experiments that included P450s and NAD(P)-binding Rossman-Fold Family Proteins, we observed the production of a highly abundant mass feature with an *m/z* value of 274.1443 in positive ionization mode, which corresponded to the expected mass ([M+H]^+^) of an oxidative coupling product that had undergone a reduction. MS/MS analysis and comparison to a standard reveled that this was in fact an intermediate in the sought-after lycorine *o-p’* pathway, norpluviine (**7**) (Figure 3, Supplementary Figure S13).

Upon deconvolution of the batch-tested candidates, it appeared that norpluviine was produced as a result of co-infiltration of *Nt*CYP96T5, a CYP96T1-like cytochrome P450 and *Nt*SDR1, a short-chain alcohol dehydrogenase/reductase. Neither *Nt*CYP96T5 nor *Nt*SDR1 alone result in the production of **7** (Figure 3C). While this suggests that, analogously to the *p-p’* and *p-o’* pathways, *Nt*CYP96T5 oxidizes 4OMN and *Nt*SDR1 then reduces the product to afford **7**, we could not detect the expected 4OMN oxidation intermediate ([M+H]^+^ = *m/z* 272.1281) when *Nt*CYP96T5 was tested alone. Instead, a set of peaks with a different mass (i.e. matching an expected formula of **7** + [O] – 2[H], [M+H]^+^ = *m/z* 288.12) was observed (Supplementary Figure S14). These unexpected oxidized mass features may be true intermediates in the biosynthesis of **7**, or could be unstable by-products that are produced by *Nt*CYP96T1 in the absence of *Nt*SDR1. We suspect that biosynthetic pathways for plant specialized metabolites are likely highly streamlined (e.g. by protein-protein interactions, or tuning of protein levels and reaction kinetics) presumably this helps prevent leakage of toxic or chemically unstable intermediates^68, 69^. By showing that detection of the expected pathway intermediate **7** required the simultaneous expression of the oxidative enzyme *Nt*CYP96T5 and the reductase *Nt*SDR1, our results highlight the importance of testing enzymes in batch as unstable intermediates that are challenging to detect using standard methods could be formed with partial pathways.

With the identification of the third CYP96T1 homolog *Nt*CYP96T5, we thus had identified P450 enzymes for all three modes of oxidative coupling and, moreover, revealed that a single set of closely related CYP96T1 homologs, perhaps formed through duplication and functional divergence, were catalyzing three different modes of oxidative couplings on the same substrate, 4OMN, to generate the basic scaffolds that give rise to the noted diversity of AmAs. Together, our results reveal a major channel to diversity generation in the Amaryllidaceae Alkaloids and lay the grounds for future engineering of these bioactive metabolites into heterologous systems.

### Site-directed mutagenesis reveals the basis for divergent regioselectivities of oxidative coupling P450s

The oxidative enzymes CYP96T1 and *Nt*CYP96T6 that generate the haemanthamine and galantamine scaffolds from the 4OMN precursor are highly similar to each other, exhibiting 91% identity at the nucleotide level (86% at the amino acid level). As gatekeeping enzymes in *p-p’* and *p-o’* scaffold biosynthesis respectively, these enzymes may be critical to determining the relative flux of 4OMN to these two bioactive natural products, galantamine and haemanthamine (Figure 3a). While the native CYP96T1 sequence produces only trace amounts of the *para-ortho’* product **2**, its homolog *Nt*CYP96T6 displays inverted regioselectivity, with the *para-ortho’* product **8** as the predominant product observed on an LC/MS Extracted Ion Chromatogram (EIC) (Figure 3D). Since galantamine, an AChE inhibitor, and haemanthamine, a eukaryotic ribosome inhibitor, appear to exhibit mutually exclusive biological activities, we sought to understand the structural basis for the divergent regioselective oxidations that produced their scaffolds. We considered that identification of the regioselectivity-determining residues in *Nt*CYP96T1 and *Nt*CYP96T6 could allow control over relative precursor flux to either of these bioactive metabolites^22, 23^.

We first looked for amino-acid differences between *Nt*CYP96T1 and *Nt*CYP96T6 sequences that resided within the substrate recognition sites (SRSs)— motifs that are known to be important for substrate recognition and binding in P450 catalysis^70^. We then searched for orthologous proteins encoded in other publicly reported transcriptome assemblies of AmA producers (namely, *Lycoris longituba* and *Narcissus pseudonarcissus*, var. King Alfred)^47, 48^ and identified residues that were conserved across orthologs. By looking for differences within the SRSs of CYP96T1 and *Nt*CYP96T6 that were consistently conserved within their respective *L. longituba* and *N. pseudonarcissus* var. King Alfred orthologs, we identified a total of four candidate sites that might impart regioselective oxidation in CYP96T1 and *Nt*CYP96T6 (Figure 3E, Supplementary Figures S15).

We next performed sequence-swapping mutagenesis for each of these four regions, and assessed the impact of swapped residues on catalytic regioselectivity. We cloned mutants for CYP96T1 that contained the corresponding *Nt*CYP96T6 residues for each of these four motifs and tested these mutants in the transient expression system in *N. benthamiana*. Of these, while three of the mutated motifs didn’t impact product regioselectivity, a single alanine on *Nt*CYP96T1, when mutated to the corresponding leucine from *Nt*CYP96T6, imparted *Nt*CYP96T1 with the ability to catalyze the *para-ortho’* oxidative coupling (Figure 3F, Supplementary Figure S16). Closer inspection of this site through sequence analysis, homology modelling and substrate docking revealed that the A308 site was present within the oxygen-binding motif of *Nt*CYP96T1, and was in close spatial proximity to the predicted binding site for its substrate 4OMN (Supplementary Figure S17).Collectively, these results reveal that alkaloid flux in daffodils can be gated by highly similar enzymes where a single amino acid mutation enables the production of completely different classes of bioactive AmAs.

### Identification of the core biosynthetic pathways for haemanthamine and galantamine

Having reconstituted the first committed steps in galantamine and haemanthamine biosynthesis in *N. benthamiana*, we next sought to identify enzymes for remaining uncharacterized steps in their biosynthesis. Since plant specialized metabolic pathways tend to co-express, we used a combination of chemical logic and co-expression analysis to identify the remaining candidate biosynthetic genes in the core haemanthamine and galantamine pathways (Figure 4, Supplementary Figure S1)^14, 71–73^.

**Figure 4:**
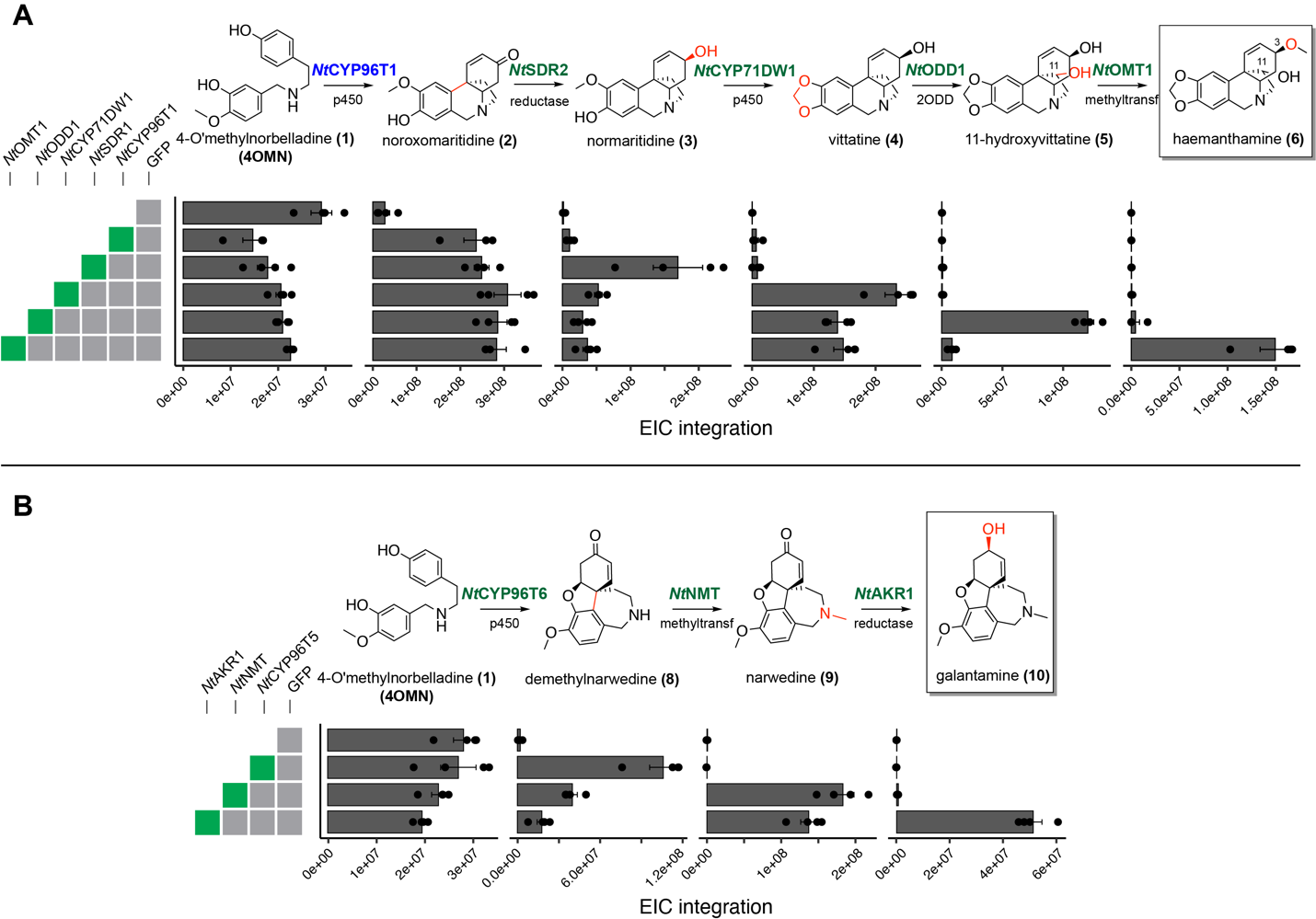
Reconstitution of the core haemanthamine (A) and galantamine (B) biosynthetic pathways in *N. benthamiana* through *Agrobacterium*-mediated transient T-DNA expression of pathway genes alongside infiltration of 200 µM of 4-O’-methylnorbelladine substrate. The gray boxes to the left indicate the genes that were co-infiltrated in *N. benthamiana* leaves. For each intermediate, the EIC peak integrations for the [M+H]^+^ ion are reported for 4 replicates. *Nt*CYP96T1 (in blue) is a paralog of *Np*CYP96T1, the *para-para’* oxidative coupling enzyme reported by Kiligore et al^33^. All other enzymes shown in green are reported herein.

Previous precursor feeding experiments have identified potential intermediates in the biosynthetic pathway to haemanthamine and galantamine^16–18^. Haemanthamine has been hypothesized to be biosynthesized through oxidative coupling of the precursor 4OMN to noroxomaritidine (**2**), followed by reduction of the ketone to normaritidine (**3**), generation of the methylenedioxy-bridge to yield vittatine (**4**), hydroxylation at the 11-position to 11-hydroxyvittatine (**5**) and finally, O-methylation to afford haemanthamine. Using the gatekeeping enzyme for *p-p’* biosynthesis (i.e. CYP96T1) as bait, we searched for co-expressing transcripts that encode enzymes that could catalyze downstream transformations in haemanthamine biosynthesis (Supplementary Figure S11).

We identified candidate keto-reductases that could reduce **2** to **3** by searching for NAD(P)-binding Rossman Fold proteins in our transcriptomic database (Methods). Through *Agrobacterium* mediated transient T-DNA expression in *N. benthamiana*, we discovered *Nt*SDR2, a cinnamoyl-CoA reductase-type enzyme that, when coexpressed with CYP96T1 (which produces **2**) and fed with the substrate 4OMN, yielded a product with the expected MS/MS fragmentation pattern for **3** (Supplemental Figure S18, Figure 4A). Next, we searched for co-expressing enzymes that could oxidize **3** to generate the methylenedioxy bridge and yield **4**. P450s are versatile oxidative enzymes that have previously been shown to catalyze methylenedioxy bridge formation from guaiacol moieties in plants^74, 75^. We therefore focused our search on candidate P450s. This led to the identification of *Nt*CYP71DW1, a CYP71-family protein that produced **4** in our transient expression system (confirmed by LC-MS/MS comparison to standard, Supplemental Figure S19, Figure 4A). We note here that stereoisomers of known AmAs including vittatine have been isolated in various Amaryllidoideae species^76^. Thus, although wherever possible, we verified the molecular identities of enzyme products against authentic standards, it is possible that the enzymes we report here may be producing metabolites that, although indistinguishable by standard reversed phase LC-MS/MS methods, may in fact be inseparable (i.e. enantiomeric) or poorly separable stereoisomers of the said compounds. For the subsequent hydroxylation of **4** to **5**, we again turned to searching for co-expressing P450s and 2-oxoglutarate dependent dioxygenases (ODDs). Both classes of enzymes belong to the largest families of metabolic enzymes in the plant kingdom, and have previously been reported to be capable of hydroxylating un-activated carbons^13, 77, 78, 79–81^. Our search led us to the identification of *Nt*ODD2, an ODD that hydroxylates **4** to yield **5** in our transient expression system, as verified by LC-MS/MS comparison to a standard. (Supplemental Figure S20, Figure 4A). For the final step in haemanthamine biosynthesis, i.e. the *O*-methylation of **5** at the C-3 hydroxyl, we searched our candidate list for coexpressing methyltransferases (MTs)^82^. Through co-infiltration in our *N. benthamiana* transient expression system alongside CYP96T1, *Nt*SDR2, *Nt*CYP71DW1, *Nt*ODD2 and the 4OMN precursor, we identified *Nt*OMT1 as 11-hydroxyvittatine O-methyltransferase that produced haemanthamine as its product (Supplementary Figures S21 and S22, Figure 4A, verified through LC-MS/MS comparison to authentic standard). Thus, with the identification of *Nt*OMT1, we discovered the core biosynthetic pathway to the eukaryotic protein synthesis inhibitor, haemanthamine, and reconstituted its production in the heterologous host, *N. benthamiana* (Figure 4A).

With the complete core pathway to haemanthamine discovered, we next turned to identifying candidate genes for the biosynthesis of the Alzheimer’s disease drug galantamine. The core pathway to galantamine requires oxidation of 4OMN to **8**, N-methylation of to **9** and reduction of the **9** ketone to galantamine^17^. Since we had already discovered *Nt*CYP96T6 as the oxidative enzyme producing **8** from 4OMN, we looked for candidate co-expressing methyltransferases for production of **9**. This led us to the identification of *Nt*NMT1, a gamma-tocopherol methyltransferase that catalyzes the production of **9** from the *Nt*CYP96T6 product **8** (Figure 4B, Supplementary Figures S23, S24 and S25). Next, to identify enzymes that could reduce **9** to galantamine, we searched for coexpressing NAD(P)-binding Rossman Fold Superfamily proteins and identified an aldo-keto reductase, *Nt*AKR1 that reduced **9** to the drug galantamine (Supplementary Figures S26, S27, S28). With the identification of *Nt*CYP96T6, *Nt*NMT1 and *Nt*AKR1, we thus identified the minimum set of core genes for the biosynthesis of the FDA-approved Alzheimer’s disease therapeutic, galantamine. Together, these newly discovered enzymes in haemanthamine and galantamine biosynthesis reveal how 4OMN is elaborated into two structurally divergent bioactive natural products with orthogonal bioactivities.

### Isotope labelling studies on Veratrum steroidal alkaloids suggest a general paradigm for defense metabolite biosynthesis in bulbous geophytes

The localization of defense toxin biosynthesis to newly developed leaf tissue in seasonal geophytes may be crucial to protecting photosynthetic tissue from herbivory^3, 83^. Having discovered that the Amaryllidacae alkaloids are produced in newly divided leaf tissue in daffodils, we sought to assess whether other non-Amaryllidaceae bulbous geophytes also exhibited a similar developmental correlate to defense toxin biosynthesis. Outside the Amaryllidaceae, several diverse plant species exhibit geophytic lifestyles with underground storage organs that support short, seasonal growth cycles. Of these, we focused our efforts on the medicinal plant genus Veratrum, which produces steroidal alkaloids like cyclopamine that is known to act as a developmental toxin through inhibition of the Sonic hedgehog signaling pathway, for which some pathway enzymes were already reported^13, 84^.

Prior feeding studies indicate that mevalonate pathway gives rise to the core terpene scaffolds in the Veratrum steroidal glycoalkaloids^85, 86^. We hypothesized that feeding ^13^C-labelled acetate, a key precursor to the mevalonate pathway, to Veratrum would result in ^13^C isotope incorporation into pathway metabolites, thus allowing us to track active biosynthesis in Veratrum tissues. Similar to our feeding experiments with daffodils, we soaked sections of *Veratrum nigrum* leaves in solutions of ^13^C-acetate and tracked the incorporation of ^13^C label into pathway metabolites (Figure 5). A targeted search for known pathway intermediates revealed incorporation of the label into a mass feature corresponding to verazine (based on the MS/MS spectra previously reported), a steroidal alkaloid intermediate known to accumulate in *Veratrum* species.^13, 87^ ^13^ (Figure 5, Supplementary Figure S30 and S31). Analysis of label incorporation into verazine across different tissues suggests that in *Veratrum*, like in daffodils, active biosynthesis in the leaf was localized to the young, newly divided zones at the leaf base. We corroborated our feeding studies with RT-PCR to track expression of a known cyclopamine pathway gene, GABAT1, relative to an actin control as internal reference, matched the ^13^C isotope labelling pattern (Figure 5). Our work sets the stage for future discovery of the biosynthetic pathways to these medicinally important family of steroidal alkaloids in *Veratrum*, and more importantly, suggests a broader developmental paradigm to defense metabolite biosynthesis in bulbous geophytes.

**Figure 5:**
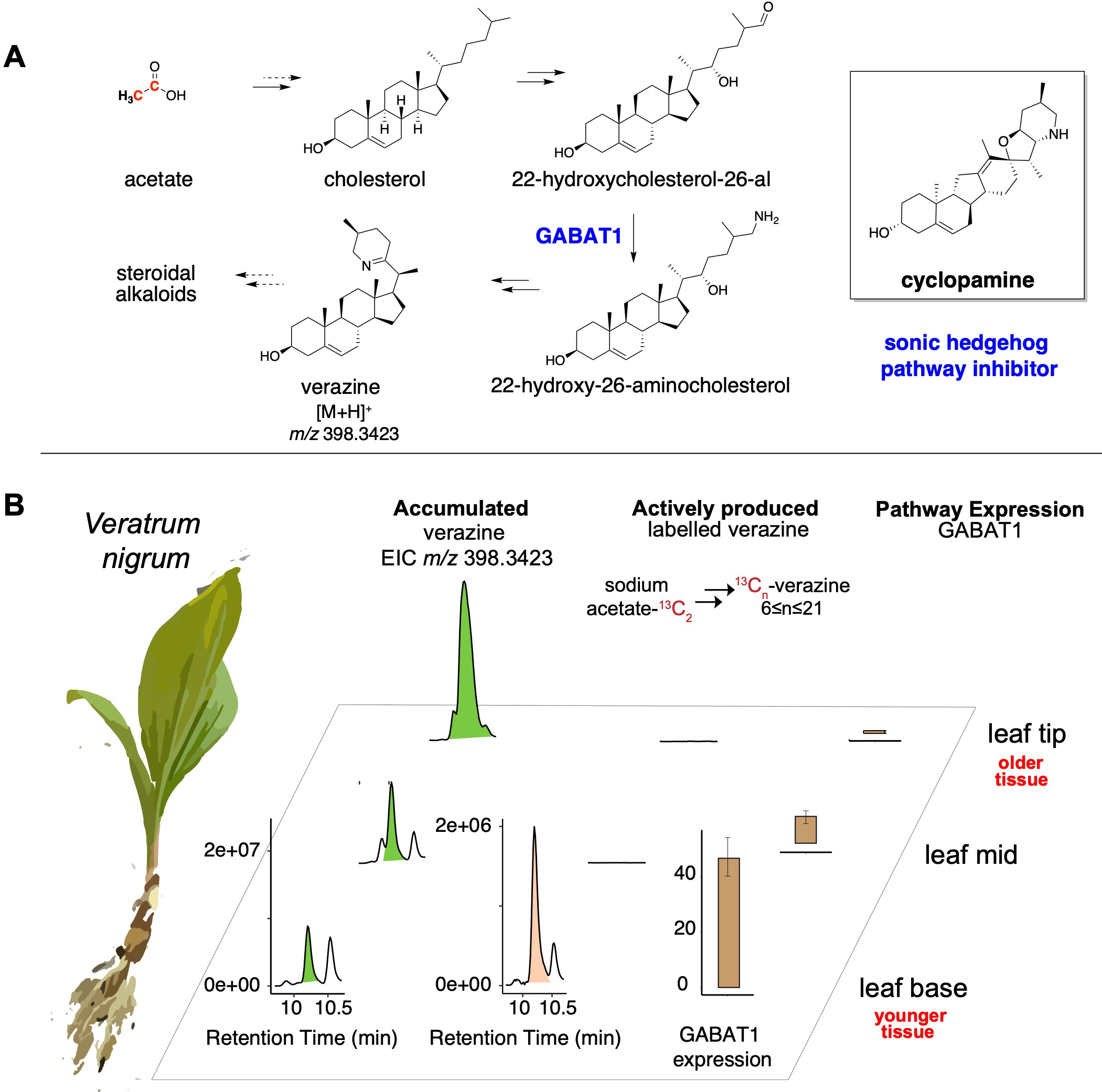
Feeding studies in *Veratrum nigrum* leaves reveal leaf bases as sites of active biosynthesis of steroidal alkaloid developmental toxins. Collectively, these data, alongside feeding studies on Amaryllidaceae alkaloids reported in Figure 2, suggest a developmental paradigm for regulation of defense metabolite biosynthesis in bulbous geophytes where young, newly developed leaf tissue is charged with defense toxins. **A)** A schematic of steroidal alkaloid biosynthesis in the genus Veratrum. Cyclopamine, a sonic hedgehog pathway inhibitor, is a steroidal alkaloid known to accumulate in plants in the Veratrum genus. Solid black arrows indicate that enzymes for the relevant chemical transformations have been reported, whereas dashed black arrows indicate that enzymes have not been reported. **B)** Feeding studies with ^13^C-labelled acetate reveal incorporation of label into verazine (N = 3 biological replicates, of which one representative replicate is shown here; unlabelled acetate was used as a control here). Extracted Ion Chromatograms (EICs) for unlabelled (natively accumulated) verazine and ^13^C labelled verazine within the same assay samples have been reported here. (The EICs for unlabelled verazine were generated by extracting for the [M+H]^+^ = *m/z* 398.3423; for the^13^C labelled verazine, EICs were extracted similarly to Supplementary Figures S30 and S31, across the envelope of ^13^C labelled verazine mass features i.e. EICs for ^13^C_n_-verzine, where 6≤n≤23). The verazine mass feature was identified solely by comparing the MS/MS fragmentation patterns of the putative verazine mass feature to a literature report, as further elaborated in Supplementary Figures S30 and S31^13^. The feeding studies were corroborated with RT-PCR expression quantification of the verazine biosynthetic enzyme GABAT1 (N=3 technical replicates)^13^. The expression fold change was quantified using the 2^-ΔΔCt^ method. All experiments measuring the expression of the pathway were performed on a separate plant.

## Discussion

As sessile organisms, plants possess an arsenal of diverse bioactive compounds to mediate their ecological interactions^88–90^. The biosynthesis of these defense metabolites is carefully coordinated and is often responsive to environmental and developmental context^91, 92^. As complex multi-cellular organisms with elaborate systems in place for transporting water and metabolites across various tissues, plants may not store defense metabolites in the same places that they produce them, making the identification of the site of active biosynthesis critical for discovering biosynthetic enzymes using expression profiling of active tissue^93, 94^.

When geophytes like daffodils release from dormancy, leaves begin to sprout from within the bulb^56, 57^. During above-ground active growth, foliage leaves photosynthesize and elongate, and flowering may occur. At the beginning of summer, these leaves wilt and translocate their photosynthate into the bulb for use as a food reserve to develop new tissue for the next growing season^95^.Here, through careful analysis and isotope-labelling studies we analyzed the patterns of metabolite variation and active biosynthesis in growing daffodil leaves. These data revealed that daffodils charge nascent, newly developed leaf tissue with toxic alkaloids as the leaf emerges above-ground from within the bulb for photosynthesis (Figure 2, Supplementary Figure S27). Thus, even though all surveyed parts of the daffodil bulb, leaves and stems contain high levels of these alkaloids, their active production is localized to only a small portion of young, growing tissue in the leaf bases within the bulb.

By diversifying a common precursor through similar, but slightly different chemical transformations, over 500+ structurally diverse metabolites are produced by plants in the Amaryllidoideae subfamily^90, 96–98^. The production of three different scaffolds from the same substrate using three slightly different versions of the same enzyme is an elegant demonstration of the parsimony underlying metabolic diversification in the Amaryllidaceae – by using duplicated homologs of the same enzyme with just small changes to their amino acid sequences, AmA biosynthesis in daffodils thus exemplifies ‘diversity oriented biosynthesis’ in the plant kingdom. Moreover, through sequence analysis and site-swapping mutagenesis on the closely related oxidative coupling enzymes *Nt*CYP96T1 (*p-p’*) and *Nt*CYP96T6 (*p-o’*), we show that the sequence space separating the differential regioselectivities of these gatekeeping enzymes in AmA biosynthesis is in fact even smaller than their 91% identities to each other suggest— a single amino acid substitution on *Nt*CYP96T1 imparted it the ability to catalyze *p-o’* oxidative coupling in addition to *p-p’* coupling. Thus, while the key end-products of the *p-p’* and *p-o’* pathways (i.e. haemanthamine and galantamine, respectively) show completely different biological activities, i.e. eukaryotic growth inhibition and AChE inhibition respectively, the enzymes that are responsible for generating their scaffolds can be different by as little as one residue. Given this low mutational barrier to switching catalytic properties, we speculate that the alterations in catalytic function may be a frequent evolutionary occurrence for these gatekeeping enzymes in AmA biosynthesis. Should such a mutation occur in nature, the cumulative alkaloid profile of the plant could be significantly altered as the precursor flux is redirected to other alkaloids, which could partly explain the high variability in alkaloid profiles observed in various AmA-producing plants^51, 99, 100^. Taken together, our work sets the stage for future discovery efforts and heterologous engineering of other classes of AmAs, including the anti-tumor compounds of the isocarbostiryl type like narciclasine^26, 101, 102^.

Finally, our work raises intriguing questions about the nature of toxin biosynthesis in other bulbous geophytes outside of the Amaryllidoieae family. Numerous additional monocotyledonous plants exhibiting geophytic lifestyles are known to produce specialized metabolites implicated in plant defense. Bulbous geophytes such as onions (*Allium cepa*) and the False hellebore (*Veratrum californicum)*, for example, exhibit very similar lifestyles to daffodils, characterized by short growing seasons followed by prolonged underground dormancy. These plants are known to produce a plethora of defense metabolites that include the various organosulfur flavor compounds and toxic steroidal glycoalkaloids, respectively^13, 107^. By uncovering a previously unknown developmental correlate to defense metabolite biosynthesis in bulbs like daffodils and Veratrum, our work suggests a paradigm for antifeedant and toxin biosynthesis in other geophytes, and lays the groundwork to the future discovery of toxin production in these important classes of plants.

### Chemical Reagents

All standard biological and chemical reagents were purchased from commercial vendors, unless noted otherwise. These include galanthamine hydrobromide (TargetMol and TCI America), lycorine (BioCrick), and 4-O’-methylnorbelladine (Toronto Research Chemicals). Standards for haemanthamine, norpluviine and vittatine were kindly provided by Dr. Lucie Cahlikova^108^. [^2^H]-4-O’-methylnorbelladine was synthesized from commercially available chemicals as previously described. ^13^C_2_-sodium acetate was purchased from Sigma-Aldrich (cat. 282014).

### Plant Materials and Growth

All experiments that were run in daffodils, except for DESI-MS imaging, were performed on blooming daffodils, *Narcissus cv. Tête-à-Tête* (*Narcissus cyclamineus × Narcissus ’Cyclataz’*) purchased from Trader Joes’ (Milgro). For DESI-MS, *Narcissus papyraceus* ‘Nir’ paperwhites were used, obtained as bulbs from Easy to Grow Bulbs (https://www.easytogrowbulbs.com/), and grown without soil over water at ambient room temperature next to a window, with the roots submerged in water. *Veratrum nigrum* plants were purchased from a commercial vendor (Edelweiss Perennials).

The *N. benthamiana* plants used for heterologous gene expression were grown in PRO MIX HP Mycorrhizae soil (Premier Tech Horticulture) at ambient laboratory temperature under growth lights with a 16/8-h light/dark cycle. 4-5 week old plants post germination were selected for *Agrobacterium-*mediated transformation.

### Stable Isotope Precursor Feeding Experiments to Track Active Metabolism

Freshly blooming daffodils were removed from soil, their roots and bulb were thoroughly washed. The outermost dry papery (tunicate) layer was removed. The basal plate, which was defined as the region with denser, darker tissue at the base of the bulb, was removed using a scalpel. Using a scalpel, successive layers of the bulb were ‘peeled’ one at a time until the innermost region of the bulb, which constituted of bases of actively growing foliage leaves, was left. The innermost bulb scale that was not a base of an actively growing foliage leaf was split into three regions – the ‘top’ region of the scale, was approximately ∼1.5 cm, and the ‘bottom’ region was approximately ∼1 cm. Each of these regions was then longitudinally split into equal sized sections, and was successively used for feeding experiments, RNA extraction and LCMS analysis of unfed tissue for assessing levels of natively accumulating metabolites. Similarly, each foliage leaf was cut transversely into eight sections of approximately 2.5-3cm each. Eight such approximately equal-sized foliage leaves were harvested from the same plant, and each leaf was used either for feeding experiments, RNA extraction or LCMS analysis.

Tissues meant for accumulated metabolite analysis & RNA extraction were immediately flash frozen in liquid nitrogen and stored at −80°C until further processing. The setup for the feeding experiments was as follows: sterile six-well plates were filled with ∼6mL of an aqueous solution of ∼405µM 4-O’-methylnorbnelladine from a 135mM d6-DMSO stock (or 3:1000 d6-DMSO in water as control). Each well contained only one section of leaf, that was then cut up into smaller pieces for maximizing direct contact with substrate solution. (Supplementary Figure S28) The covered 6-well plates were left on a lab bench for four days. Four days later, the tissue sections were moved to pre-weighed 2mL Eppendorf SafeLock tubes, flash frozen, and lyophilized. Lyophilized tissue was homogenized on a ball mill (Retsch MM 400) using 5-mm diameter stainless steel beads, with shaking at 25Hz for 2 min. 35µL of 80% methanol in water was used per mg dry weight of plant tissue to extract metabolites from plant tissue. Extracted samples were vortexed and kept at room temperature for 30 min, and then centrifuged at 16,000 x g for 5 min to pellet plant debris. The supernatant was filtered through 0.45 µm PTFE filter plates before analysis by LC-MS. RT-PCR experiments were used to corroborate the results from feeding studies, with oligos designed based on methods described before^47^ (Supplementary Table S4).

All feeding experiments were performed on leaves from the same plant. Like described above for daffodils, the outermost tunicate layers of the bulbous tissue of a *Veratrum nigrum* plant were removed. Subsequently, the layers of the bulb were successively peeled until the leaf bases were exposed. The bottommost ∼1cm of the leaf base was isolated as the leaf base. A ∼1 cm section of the leaf protruding out from the neck of the bulb was isolated as “leaf mid” and the topmost ∼1cm section of the leaf was isolated as the “leaf top.” Small pieces of these sections were made using scissors, and sections were immediately immersed in 6-well plates containing substrate solution (10mM ^13^C_2_-sodium acetate) or water (or, unlabelled 10mM sodium acetate, as reported in Figure 5). The feeding assays were run for 6 days. Metabolite extracts were generated in the same manner as described above for daffodils (80% methanol extraction). On a separate *Veratrum nigrum* plant, RNA extraction and cDNA synthesis was performed on leaf base, mid and tip sections (similarly to daffodils, see methods, RNA extraction) and RT-PCR assays were performed to quantify expression of the known verazine pathway gene GABAT1 using Actin as an internal reference, based on the Actin target sequence previously reported (Supplementary Table 4)^109^.

### RNA extraction

Daffodil tissues frozen at −80 °C were homogenized to a fine powder under liquid nitrogen using a mortar/pestle that were pre-treated with RNAse Zap RNAse Decontamination Solution (Thermo Fisher Scientific). Two separate RNA extraction protocols were used for different daffodil tissues: for root, flower and stem tissue, RNA was extracted from ground tissue using the RNEasy Plant Mini Kit (Qiagen), with DNAse on column digestion. For leaf and bulb scale tissue, RNA was extracted using TRIzol reagemt (Thermo Fisher Scientific) and cleaned up using the RNEasy Plant Mini Kit (Qiagen) with DNAse on column digestion. RNA quantification for each sample was performed via NanoDrop and Qubit HS (high sensitivity) RNA assay kit (Thermo Fisher Scientific), and quality was further assessed by measuring size distribution using an Agilent 2100 Bioanalyzer.

### Preparation and sequencing of Illumina HiSeq 4000 libraries

Library preparation for Illumina NovaSeq 6000 S4 PE150 sequencing was performed using the NEBNext® Ultra™ II Directional RNA Library Prep Kit for Illumina® (New England Biolabs), as per the manufacturer instructions. Individual libraries were barcoded with an internal index set at Novogene. Inserts were size selected with NEBNext beads to have an average insert size of 300-600 bp, and the quality and size distribution of each library was confirmed with analysis using a High Sensitivity DNA chip on a 2100 Bioanalyzer (Agilent). Equimolar amounts of libraries for each sample (28 total) were pooled for paired-end sequencing within a single Novaseq 6000 lane (S4 PE150 sequencing).

The quality of raw Illumina reads was assessed using FastQC (Babraham Bioinformatics, Babraham Institute), and reads were processed to remove adaptors using Trim Galore < --paired --retain_unpaired --length 40 > and then were subject to quality trimming using Trimmomatic^110^ <LEADING:5 TRAILING:5 SLIDINGWINDOW:5:10 MINLEN:50>. Following trimming, reads were again assessed using FastQC to confirm quality. The quality of the transcriptome was also assessed using bowtie2 mapping of the Illumina short reads to the PacBio Iso-Seq transcriptome.^111^

While these processed reads were used for quantification against the PacBio Iso-Seq transcriptome, we also generated a *de novo* transcriptome assembly using Trinity^112^ from Illumina reads to capture the higher sequencing depth and broader range of tissues sequenced through Illumina sequencing. Illumina *de novo* transcriptome assembly quality was assessed using BUSCO for completeness (86.6% complete BUSCOs) and redundant contigs were removed using CD-HIT EST with a 98% identity threshold, which yielded 1,411,754 contigs. Identifiers for RNA-seq tissue samples are listed in Supplementary Table S1.

### Preparation and Sequencing of PacBio Iso-Seq libraries

Sequencing libraries were prepared as previously described.^53^ For PacBio Iso-Seq, only a subset of tissues was sequenced – namely F_2 (inflorescence), SB_1 (stem base), LT_6 (leaf top), LB_2 (leaf bae) and B_20 (bulb scale). Preparation and sequencing of the PacBio SMRT sequencing library was performed by QB3 Genomics, UC Berkeley, Berkeley, CA, RRID:SCR_022170 on the PacBio Sequel II System instrument using Sequel II 2.1 binding kit with a movie time of 30 hours. RNA concentration was normalized to < 5 ng/µL using the Qubit HS RNA Kit, and then capillary electrophoresis was performed using the Agilent Bioanalyzer 2100 instrument, using the Eukaryotic Total RNA Pico Assay, following manufacturer’s instructions, with a 2 min denaturation at 70°C followed by a 2 min cold-shock on ice in a freezer. Preparation of the PacBio SMRT sequencing library was performed as described in the Iso-Seq^TM^ Template Preparation for Sequel® Systems manual using the NEBNext® Single Cell/Low Input cDNA Synthesis & Amplification and Pacific Biosciences Iso-Seq Express Oligo Kit with barcoded primers listed in the manufacturer supplied protocol to perform first-strand cDNA synthesis (standard purification) and PCR amplification. Barcoded cDNA from different samples was pooled in equimolar quantities and a single library was prepared from this pooled cDNA using the SMRTbell Express Template Prep Kit 2.0 following the standard protocol. Raw data from this experiment were processed using the default parameters for the Iso-Seq 2 pipeline available via SMRT® link software (PacBio, https://github.com/PacificBiosciences/IsoSeq/blob/master/isoseq-clustering.md). This processing generated a final sequence file of 432,086 high quality transcript isoforms. We then used CD-HIT-EST (2) with a cut-off at 97% identity to remove potentially redundant transcript sequences, which resulted in a final total of 90,359 unique transcripts. The quality of the assembly was assessed by mapping short read Illumina sequencing reads (see Methods) to the long read transcriptome assembly (91.2% overall alignment rate) and BUSCO (89.1% completeness).

### Analysis of RNA-seq Data

Peptide sequences were predicted from the PacBio generated transcriptome using TransDecoder^113^. The longest ORFs for each transcript were annotated with best-hit Pfam terms and BLASTp to the *Arabidopsis proteome* (Araport11).^114, 115^ Quantification of processed Illumina reads against the PacBio Iso-Seq Transcriptome was performed using Kallisto.^116^ For all downstream expression analysis, TMM-normalized CPM expression values were used.

### cDNA synthesis for cloning

cDNA was prepared from extracted RNA (see Methods) using the Superscript IV First Strand Synthesis Kit.

### Cloning of candidate biosynthetic genes

Candidate biosynthetic genes were cloned from cDNA generated from leaf bases or homogenized bulbs from *N. pseudonarcissus* RNA extracted as described above. cDNA was prepared from extracted RNA using the Superscript IV First Strand Synthesis Kit. Coding sequences were amplified via PCR using Q5® High Fidelity DNA Polymerase (New England Biolabs) as per the manufacturer instructions. Oligonucleotide primers were designed to target the 5’ and 3’ ends of the coding sequence for each gene, and each primer was synthesized such that the targeted amplicon had 5’ and 3’ homologous regions for subsequent Gibson assembly into the appropriate expression plasmid. Following PCR amplification, amplicon products were assessed via gel electrophoresis on a 1% (w/v) agarose gel, and PCR product was either directly used for construction of the transient expression construct or was, where needed, gel extracted and purified using the Zymoclean Gel DNA Recovery Kit (Zymo Research).

For assembly of the transient expression construct, pEAQ-HT was digested with AgeI and XhoI restriction enzymes. PCR amplicons were inserted into digested pEAQ-HT using the NEBuilder® HiFi DNA Assembly Mix (New England Biolabs), as per the manufacturer instructions. These reactions were then directly transformed into either NEB 5α or NEB 10ß *Escherichia coli* cells (New England Biolabs), which were grown on Luria-Bertani (LB) agar plates (50 μg/mL kanamycin) at 37 °C overnight to select for positive transformants. Successful insertion of the desired gene was confirmed by screening with colony PCR, and positive colonies were then used to inoculate 5 mL LB liquid cultures (50 μg/mL kanamycin), which were grown overnight on a culture rotary drum at 37 °C. Plasmid DNA was then purified from these cultures using the ZR Plasmid Miniprep Kit (Zymo Research). Inserts from isolated plasmids were then Sanger sequenced to confirm the identity and correct sequence of the insert (ELIM Biopharm Inc.).

pEAQ-HT expression constructs were transformed into *Agrobacterium tumefaciens* (GV3101:pMP90) using the freeze-thaw method, and transformed cultures were plated on LB agar (50 μg/mL kanamycin and 30 μg/mL gentamycin) and grown at 30 °C for two days to select for positive transformants, which were verified by colony PCR using *Taq* Polymerase (NEB). Positive colonies were used to inoculate 5 mL liquid LB cultures (30µg/mL gentamycin and 50µg/mL kanamycin) which were grown for two days on a culture rotary drum at 30 °C and then used to make 25% glycerol stocks that were snap frozen in liquid nitrogen and stored at −80°C for future use.

### *Agrobacterium*-mediated transient expression

Transient expression assays in N*. benthamiana* were performed as previously described.^14^ In brief, Glycerol stocks of *Agrobacterium tumefaciens* strains (GV3101:pMP90) containing the relevant pEAQ-HT expression construct were either streaked onto an LB agar plate (supplemented with 50 μg/mL kanamycin and 30 μg/mL gentamycin) and grown at 30°C for two days, or were alternatively used to inoculate 7mL liquid LB cultures (supplemented with 50 μg/mL kanamycin and 30 μg/mL gentamycin) that were then grown at 30°C for two days. Bacterial lawns were collected from the plates using a 1 mL pipette tip or from liquid cultures by centrifugation at 4,000 x g and resuspended in 1 mL of *Agrobacterium* induction medium (10 mM MES, pH = 5.6, 10 mM MgCl_2_, 150 μM acetosyringone), and centrifuged at 5,000 x g for 5 minutes to pellet cells. The pellet was resuspended in 1mL of fresh *Agrobacterium* induction medium. This resuspension of cells was incubated at room temperature for 2-3 hours, after which the optical density of the cells (OD_600_) was measured for each strain. Each strain was then diluted into induction medium to an OD_600_ of 0.3 for infiltration, such that in infiltration experiments where multiple strains were co-infiltrated into a single leaf, each individual strain was at a final OD_600_ of 0.3. These individual strains or mixtures of strains were then infiltrated into the abaxial side of 4-5 week-old *Nicotiana benthamiana* leaves using a 1 mL needleless syringe. Each combination of strains being tested was infiltrated into three leaves, such that each leaf belonged to a different plant. After 3-4 days of infiltration, 200 µM of 4-O’methylnorbelladine substrate was infiltrated into the same leaves. 1-2 days after infiltration of substrate, ∼1 square inch of infiltrated leaves was excised using scissors and placed in pre-weighed 2 mL Safe-lock tube (Eppendorf) which were snap frozen in liquid nitrogen, and lyophilized to dryness for subsequent metabolite extraction.

### LC-MS analysis of metabolites

Samples were analysed by reversed-phase liquid chromatography using a Gemini NX-C18 column (Phenomenex, 5 μm, 2 mm × 100 mm) with an injection volume of 2µL unless otherwise noted, and a mobile phase, unless otherwise noted, consisting of two buffers: water with 0.1% ammonium hydroxide solution (buffer A) and 95% acetonitrile, 5% water with 0.1% ammonium hydroxide solution (buffer B).

Extracts from *N. pseudonarcissus* feeding studies (Figure 2, Supplementary Figures S4, S5) were analysed on an Agilent 1290 HPLC (high-performance liquid chromatography) coupled to an Agilent 6545 quadrupole time-of-flight (qTOF) mass spectrometer with an electrospray ion source (dual-inlet Agilent Jet Steam) that was used to collect MS data in positive ion mode (parameters: mass range: *m/z* 100-1700; drying gas: 300°C, 12 L/min; nebulizer: 10 psig; capillary: 3500 V; fragmentor: 100 V; skimmer: 50 V; OCT 1 RF Vpp: 750 V; 2.50 spectra per second). The first minute of each run was discarded to avoid salt contamination of the MS apparatus. A flow rate of 0.4 mL/min was used with the following 26 min gradient method: 0-1 min, 3% B; 1-16 min, 3-50% B; 16-17 min, 50-97% B; 17-22 min, 97% B; 22-23 min, 97-3% B; 23-26 min, 3% B.

All metabolite extracts from transient infiltration assays, except the characterization of CYP96T1-like enzymes’ regioselectivities and their site-directed mutagenesis assays, (Figure 4, Supplementary Figures S13, S14, S18, S19 S20, S22, S25, S26, S27, S28) were analysed on an Agilent 1290 HPLC instrument coupled to an Agilent 6546 quadrupole time-of-flight (qTOF) mass spectrometer with an electrospray ion source (dual-inlet Agilent Jet Steam) that was used to collect MS data in positive ion mode (parameters: mass range: *m/z* 100-1700; drying gas: 400°C, 12 L/min; nebulizer: 20 psig; capillary: 4000 V; fragmentor: 180 V; skimmer: 45 V; OCT 1 RF Vpp: 750 V; 1 spectra per second). The first half minute of each run was discarded to avoid salt contamination of the MS apparatus. A flow rate of 0.4 mL/min was used with the following 22 min gradient method: 0-1 min, 3% B; 1-16 min, 3-50% B; 16-17 min, 50-97% B; 17-19.5 min, 97% B; 19.5-20 min, 97-3% B; 20-22 min, 3% B.

All metabolite extracts for the characterization of CYP96T1-like enzyme regioselectivities and their site-directed mutagenesis (Figure 3c,3d,3f, Supplementary Figure S16) were analysed on an Agilent 1260 HPLC with an injection volume of 5µL coupled to an Agilent 6520 quadrupole time-of-flight (qTOF) mass spectrometer with an electrospray ion source that was used to collect MS data in positive ion mode (parameters: mass range: *m/z* 100-1700; drying gas: 325°C, 10 L/min; nebulizer: 35 psig; capillary: 3500 V; fragmentor: 150 V; skimmer: 65 V; OCT 1 RF Vpp: 750 V; 1.41 spectra per second). The first half minute of each run was discarded to avoid salt contamination of the MS apparatus. A flow rate of 0.4 mL/min was used with the following 29 min gradient method: 0-1 min, 3% B; 1-16 min, 3-50% B; 16-17 min, 50-97% B; 17-22 min, 97% B; 22-23 min, 97-3% B; 23-29 min, 3% B.

All metabolite extracts from *Veratrum nigrum*, (Figure 5, Supplementary Figures S30 and S31) were analysed on an Agilent 1290 HPLC instrument coupled to an Agilent 6546 quadrupole time-of-flight (qTOF) mass spectrometer with an electrospray ion source (dual-inlet Agilent Jet Steam) that was used to collect MS data in positive ion mode (parameters: mass range: *m/z* 100-1700; drying gas: 400°C, 12 L/min; nebulizer: 20 psig; capillary: 4000 V; fragmentor: 180 V; skimmer: 45 V; OCT 1 RF Vpp: 750 V; 1 spectra per second). The first half minute of each run was discarded to avoid salt contamination of the MS apparatus. A flow rate of 0.4 mL/min was used with the following 22 min gradient method: 0-1 min, 3% B; 1-16 min, 3-50% B; 16-17 min, 50-97% B; 17-19 min, 97% B; 19-20 min, 97-3% B; 20-21 min, 3% B. The mobile phase used was A: water with 0.1% formic acid and B: acetonitrile with 0.1% formic acid.

### Synthesis of stably labelled [^2^H]-4-O’-methylnorbelladine for feeding studies

[^2^H]-4-O’-methylnorbelladine was synthesized from tyramine and isovanillin as previously described, with modifications.^33^ 1096 mg (8 mmol) of tyramine and 1216 mg (8 mmol) of isovanillin were mixed together in 40 mL of methanol and stirred for 12-16 hours at room temperature. Sodium borodeuteride (Sigma Aldrich) i.e. NaBD_4_ used as reducing agent to provide the benzylic deuterium. Subsequently, NaBD_4_ was added slowly over an ice-cooled reaction mixture. The reaction mixture was evaporated using a rotary evaporator, and resuspended in DMSO and directly loaded onto a BioTage c18 Sfar 12 g column for compound isolation, using acetonitrile with 0.1% formic acid and water with 0.1% formic acid as the mobile phases. Elution of the target compound was tracked in parallel by UV absorbance (280nm) and LC/MS. Eluting fractions containing the target compound were frozen in liquid nitrogen and lyophilized to dryness. Chemical identity of the product was confirmed by ^1^H-NMR (acquired using a Varian Inova 600 MHz spectrometer at room temperature in DMSO-d_6_, Supplementary Figure S32) and the purity of the product was assessed using LC/MS and the [^2^H]-4-O’-methylnorbelladine concentration was estimated against a standard curve of commercially available, unlabeled, 4-O’-methylnorbelladine standard, by calculating peak areas for different concentrations of the standard.

### Metabolite extraction

Samples lyophilized in pre-weighed tubes were weighed to calculate dry mass of material. These samples were homogenized to a fine powder in 2 mL Safe-Lock tubes (Eppendorf) using 5-mm stainless steel beads on a ball mill (Retsch MM 400) with shaking at 25 Hz for 2 min. To extract metabolites from homogenized samples a solution of 80% methanol 20% water (*v/v*) solution was used, with 25 or 35µL per milligram of tissue dry weight for *Narcissus* tissues, or 35µL per milligram for *N*. *benthamiana* leaves. Samples were vigorously vortexed upon addition of the extraction solution and were incubated at room temperature for 30 min. Samples were then centrifuged at 16,000 x g for 5 min to pellet debris, and were subsequently filtered through a 0.45 µm PTFE Filter Plate (MultiScreen Solvinert, hydrophilic, PTFE, Fisher) with a collection plate, sealed with Silicone Sealing Mats (Agilent) and used for LC-MS analysis.

### Metabolomic Analyses

LC-MS data were analysed using Agilent MassHunter Software. Peaks were extracted for target masses using a 20-100 ppm mass tolerance, and the area under the peak was calculated using Agilent MassHunter Quantitative Analysis Software. Peaks integrated by the Quantitative Analysis Software were manually inspected to ensure correct peak selection and integration.

### Site Directed Mutagenesis

To identify the substrate recognition sites on the candidate P450s, protein alignment was performed using Clustal Omega in Geneious (default parameters) with *St*CYP86A33, for which the substrate recognition sites (SRSs) have been reported.^117^ The resulting alignments were used to define the boundaries of the SRSs on P450s of interest. CYP96T1 and CYP22 homologs were identified in the publically available transcripomes of *Lycoris longituba*^48^ and *N. pseudonarcissus* var. King Alfred^47^ using reciprocal best blast (nucleotide), and where more than one candidate homologs were found, all hits with >95% BLASTnt identity were designated as homologs. For CYP22, only one partial transcript was found in the King Alfred transcriptome, and this partial transcript was included in the analysis. Open Reading Frame (ORF) predictions were used to generate putative protein sequences from the CYP96T1-like and CYP22-like sequences. These sequences were then aligned in Geneious using Clustal omega with the default parameters. To identify candidate residues important for regioselectivity, conserved differences between CYP96T1-like sequences and CYP22-like sequences were identified. This led to the identification of a total of 6 residues, corresponding to three different SRSs. We individually mutated the four candidate regioselectivity-imparting sites on CYP96T1 to the corresponding residues on the CYP22 sequences. The mutants were generated using a combination of PCR/Gibson assembly or were directly synthesized as gene fragments. Oligos and templates used for site-directed mutagenesis experiments are listed in Supplementary Tables S2 and S3.

### Homology Modelling

SWISS-Model and SwissDock were used with <u>6l8h.1.A</u> (Carotene epsilon-monooxygenase, chloroplastic) as a template and 4-O’-methylnorbelladine **<u>ZINC1713519</u>** as a substrate.^118, 119^ Then, Chimera (UCSF) was used to visualize docking configurations. Models with low FullFitness docking configuration that were within the hydrophobic core of the enzyme were chosen for further examination in PyMol.

### Mass spec imaging sample prep

*Narcissus papyraceus* ‘Nir’ (i.e. paperwhite) plants were used for Mass Spec imaging analysis, due to changes in seasonal availability of *Narcissus* cv. Tête-à-Tête plants. Leaves were obtained from growing paperwhites that had at least 2-3 inches of foliage emerged above the neck of the bulb. Each leaf was cut into two vertical sections – the top half and the bottom half. Each of these halves was directly immersed into a 7mL solution of [^2^H]-4-O’-methylnorbelladine (∼405 µM) (or 3:1000 DMSO in water as control) in a 50mL Falcon Tube. After 5-7 days, the basal ∼1 inch of each of the leaf halves were embedded in Optimal Cutting Temperature Compound (OCT) in Peel-A-Way embedding molds (Sigmma-Aldrich) and frozen in a slurry of dry ice and ethanol. These embedded molds were sectioned on a Leica 3050 S Cryostat, with 35µm sectioning thickness, and transferred to Microscope Slides (VWR, Superfrost® Plus). Leaf transverse sections tended to lose structural integrity at room temperature, and so the slides were immediately stored at −80°C, and were lyophilized for 20-30min to dryness for preventing delocalization of metabolites within the tissue. The slides were imaged on an EVOS™ FL Auto 2 Imaging System (Thermo Scientific™ Invitrogen™).

### Desorption electrospray ionization Mass Spectrometry (DESI-MS) Acquisition and Analysis

A commercial DESI source (Prosolia Inc., Indianapolis, IN, US) was coupled with the orbitrap mass spectrometer (Velos Pro, Thermo Fisher, MA, US) to conduct mass spectrometry imaging experiments. DMF-ACN (1:1, v/v) was used as the spray solvent with a flow rate at 0.75 μL/min, and a nebulizer gas pressure at 125 psi. The impact angle between the DESI sprayer and the moving stage was set at 56°. A high voltage of 5.0 kV was applied onto the sprayer to generate the charged microdroplets to release analytes from the section. The MS inlet capillary temperature was set at 300 °C. Positive full MS scans were employed over the range of *m/z* 250-320. The automatic gain control (AGC) was set to the off position to keep the scan rate constant. Omni Spray 2D software (Prosolia, Inc., Indianapolis, IN, USA) was used to control the 2D moving stage and to trigger MS data acquisition. The construction of ion images was achieved using a self-coded program run in MATLAB (Mathworks, Natick, MA, USA). The identity of displayed ions was verified through tandem MS/MS (35 V collision energy, CID mode with resolution set to 120000)

## Data Availability

RNA-seq data have been deposited in the National Center for Biotechnology Information (NCBI) Sequence Read Archive (SRA) database under the BioProject ID XXXXX. Cloned sequences have been deposited in GenBank (accession nos. XXXX). All other raw/source data are available upon request.

## Acknowledgements

This work was supported by National Institutes of Health (NIH) R01 grant GM121527. We are grateful to Prof. L. Cahlíková (Charles University in Prague) for providing authentic standards for several alkaloids, Dr. D. Wegnier (Stanford University) for insightful discussions on plant biology, Dr. A.-M. Faust (Novartis Institute of Biomedical Research) for early project discussions, J. Liu and Dr. K. Smith for assistance with NMR analysis.

## Author Contributions

N.M. and E.S. contributed to the study design. N.M. performed the research, Y.M. performed data collection for Figure 2C. N.M., Y.M., R.K., R.Z., and E.S. analysed and interpreted data. N.M. and E.S. wrote the manuscript with input from the other authors.

