## Supplementary Materials for "A developmental gradient reveals biosynthetic pathways to eukaryotic toxins in monocot geophytes"

**Supplementary Figures and Tables**

**Supplementary Figures**

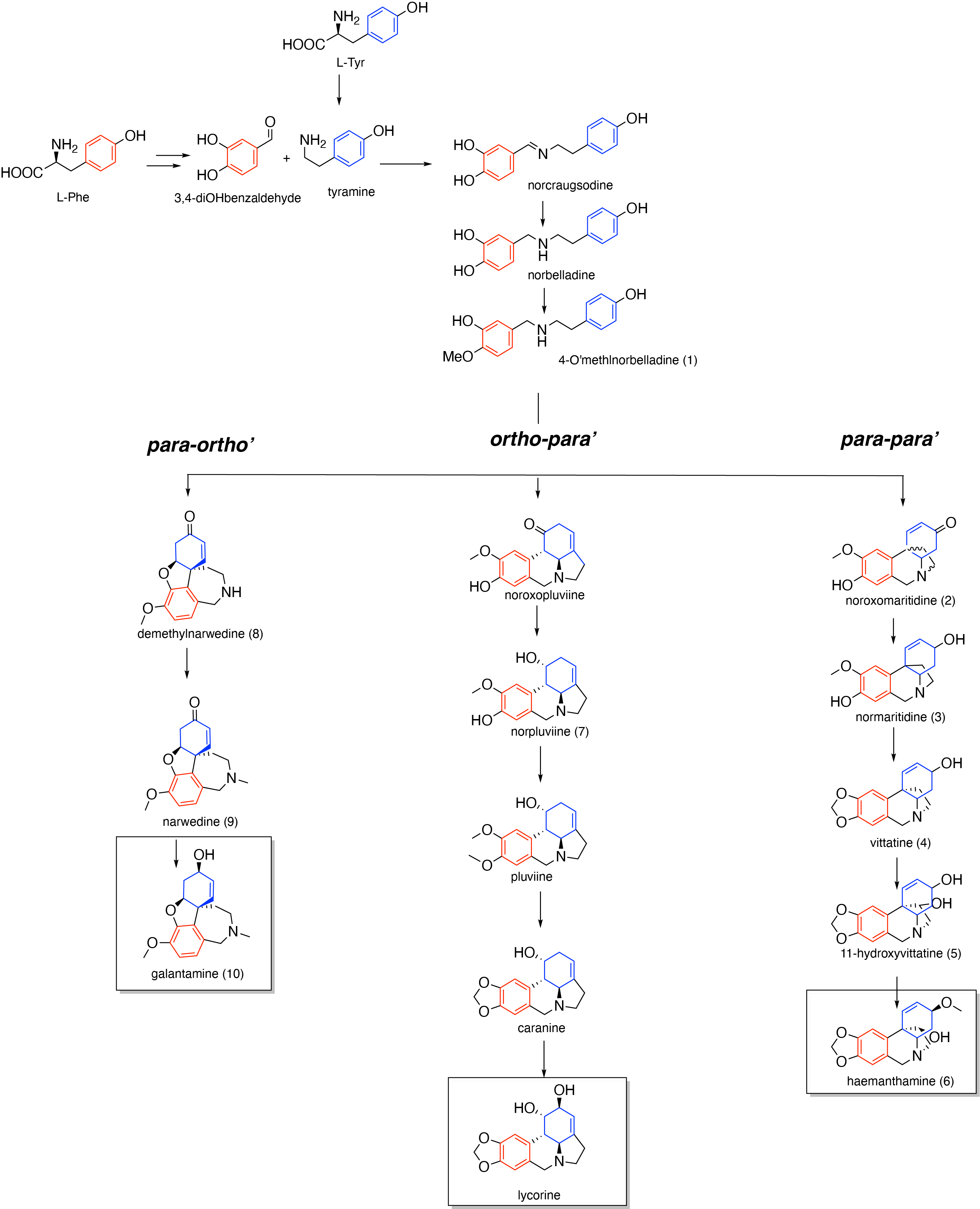

**Supplementary Figure S1:** The core (proposed) biosynthetic pathways to the key bioactive Amaryllidaceae alkaloids, galantamine *(para-ortho’)*, lycorine *(ortho-para’)* and haemanthamine (*para-para’*).

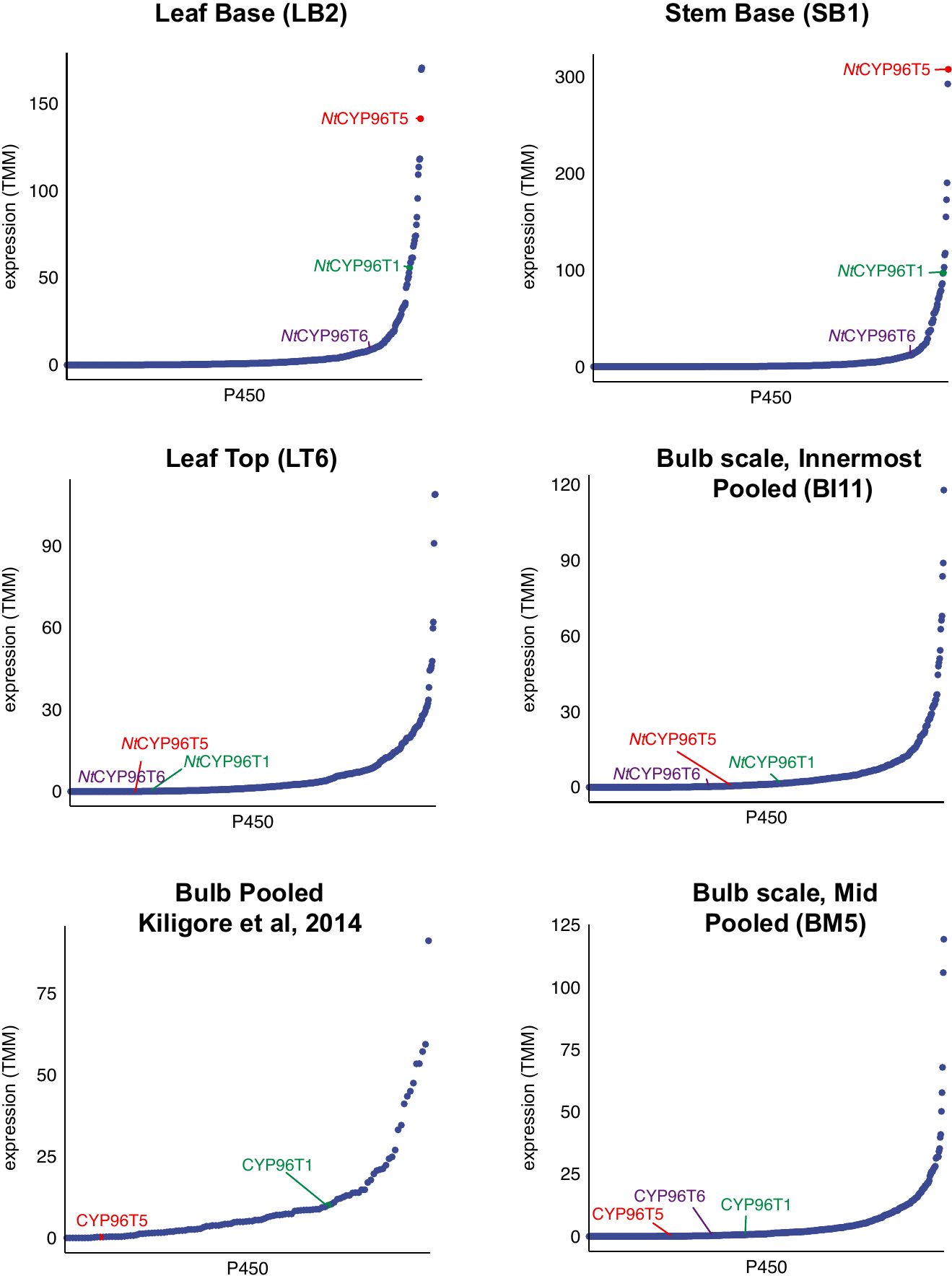

**Supplementary Figure S2:** The expression of P450s in different tissues sequenced via RNA-seq. The expression of oxidative coupling pathway enzymes (i.e. *Nt*CYP96T1, *Nt*CYP96T5 and *Nt*CYP96T6) has been labelled. For comparison, the expression of the respective enzyme best-BLAST homologs in a publicly available daffodil bulb transcriptome has been also plotted. The identification of the site of active biosynthesis enabled us to appropriately sample the site of active biosynthesis– the pathway enzymes *Nt*CYP96T1, *Nt*CYP96T5 and *Nt*CYP96T6 are amongst the most highly expressed P450s in leaf and stem bases.

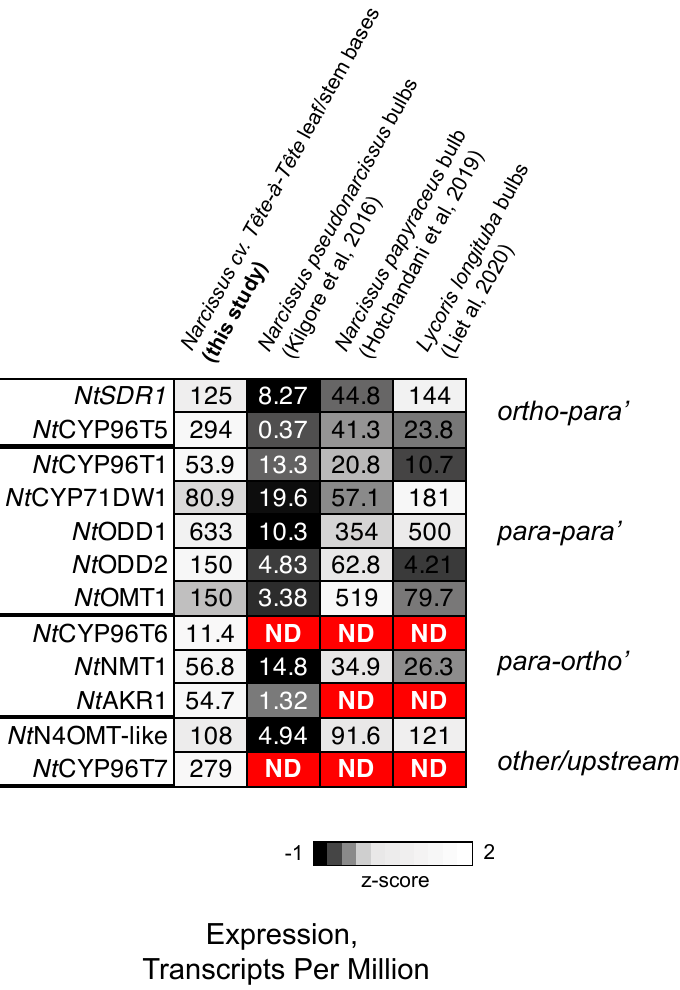

**Supplementary Figure S3**: Expression of pathway genes/ their best-BLAST homologs in publicly available transcriptomes for AmA producers. The values in the boxes denote the expression of homologs of pathway genes in Transcripts Per Million (TPM). Boxes are coloured to highlight transcriptomes with the highest expression levels for pathway genes

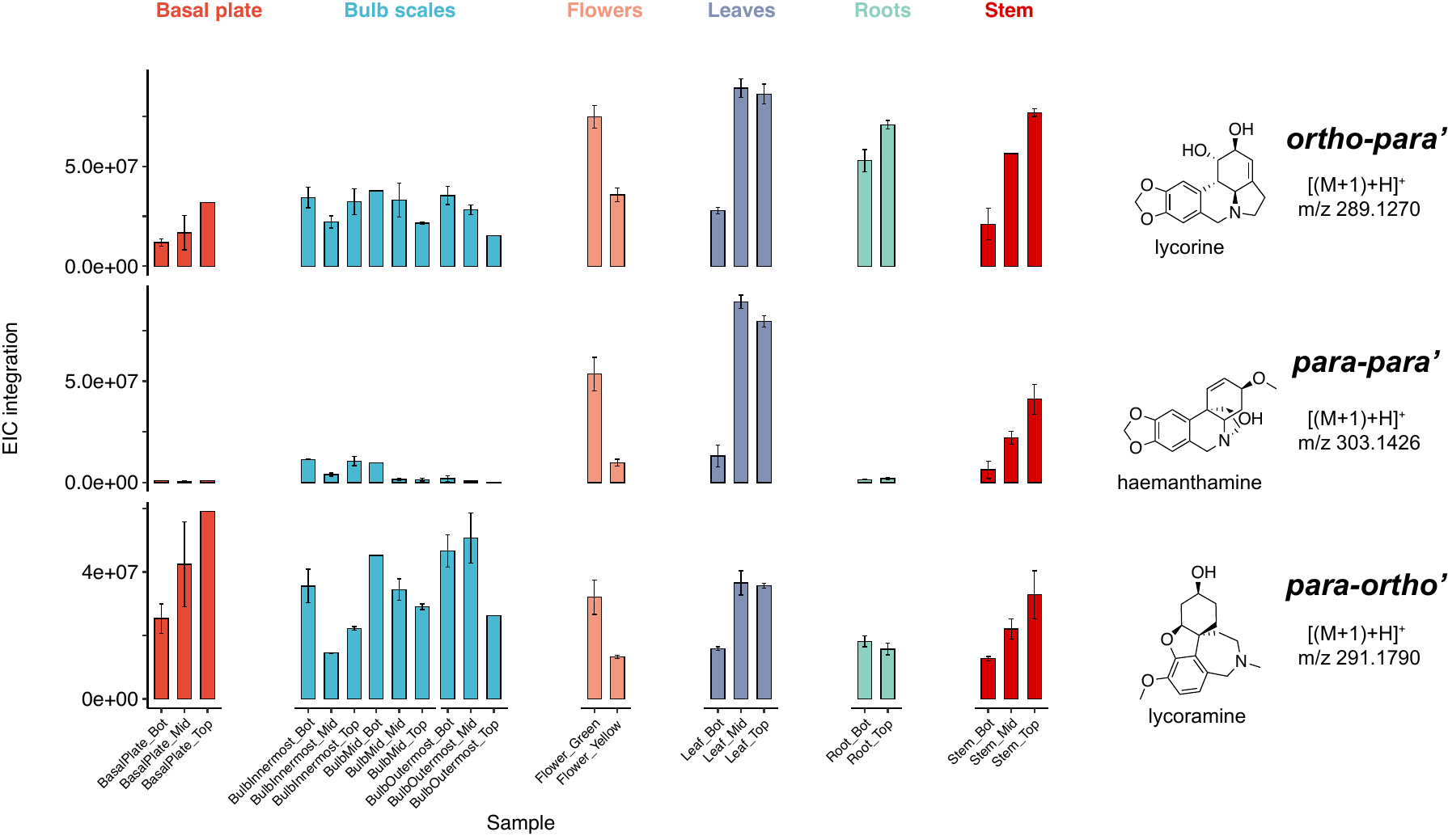

**Supplementary Figure S4:** The accumulation of key alkaloids of each, p-p, o-p and p-o classes was tracked through LC-ESI-MS in a single *Narcissus* cv. Tête-à-Tête plant at full-bloom. N=2 or 3 for each tissue. The abundances of the alkaloids are generally highest in flowers, roots and stems.

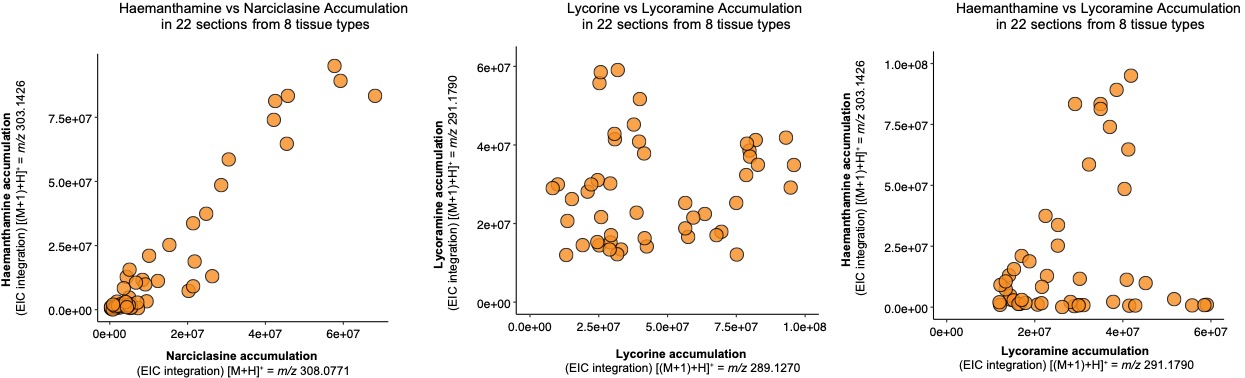

**Supplementary Figure S5:** The accumulation of key alkaloids of each, p-p, o-p and p-o classes was tracked through LC-ESI-MS in a single *Narcissus* cv. Tête-à-Tête plant at full-bloom in 22 different samples that were derived from eight different tissues shown in Figure S4. While alkaloid levels for the *para-para’* pathway metabolites haemanthamine and narciclasine appear correlated across different tissues, the levels of metabolites across different branches of the pathway (i.e. lycorine is o-p and lycoramine is p-o) appear not as well correlated across all tissues.

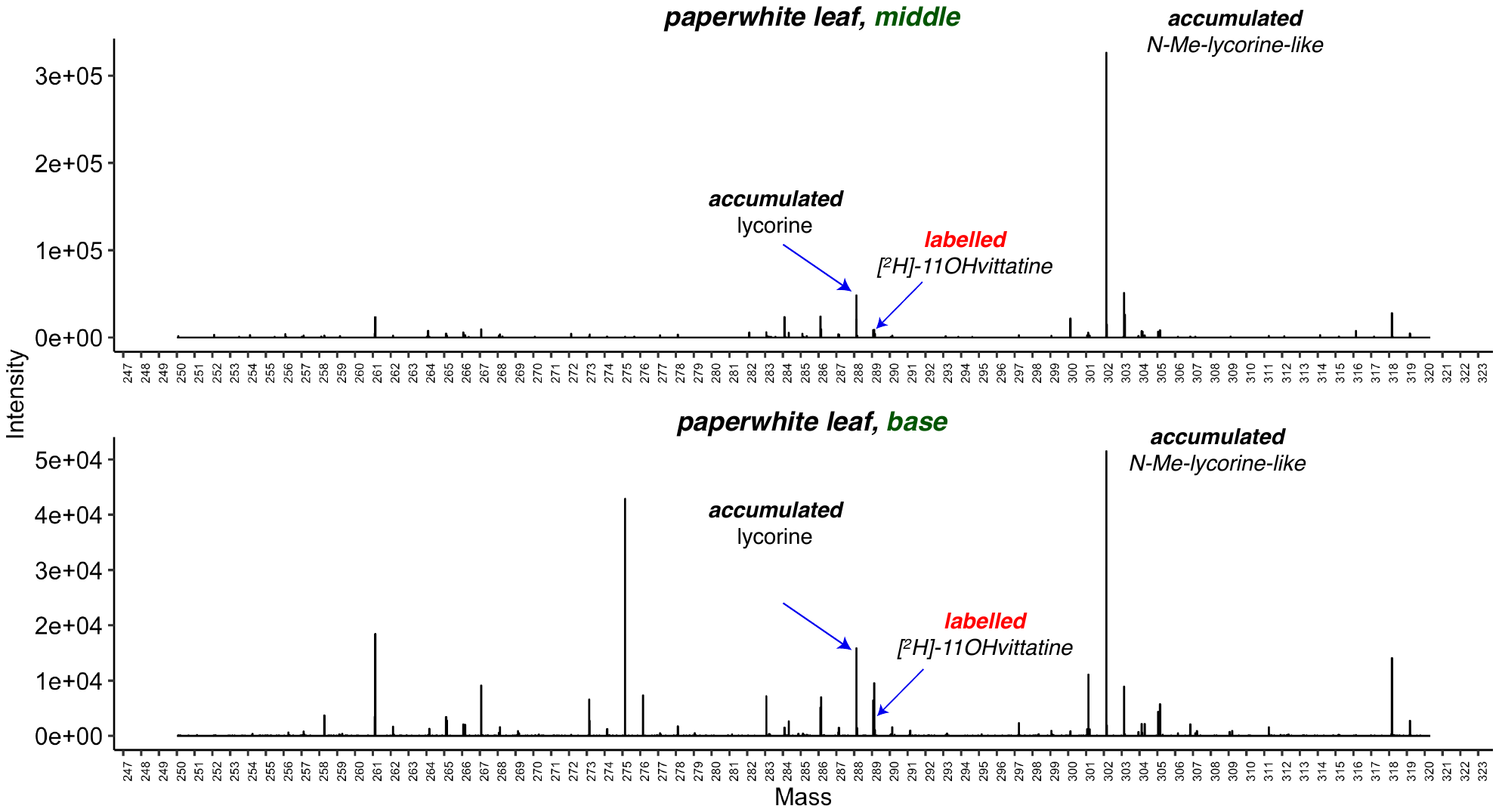

**Supplementary Figure S6:** DESI-MS spectra of [^2^H]-4-O’-methylnorbelladine fed *N. papyraceus* (paperwhite) leaf base and mid- sections. These data confirm that greater abundances of label are detected in the younger leaf bases relative to the older, middle section of the leaf.

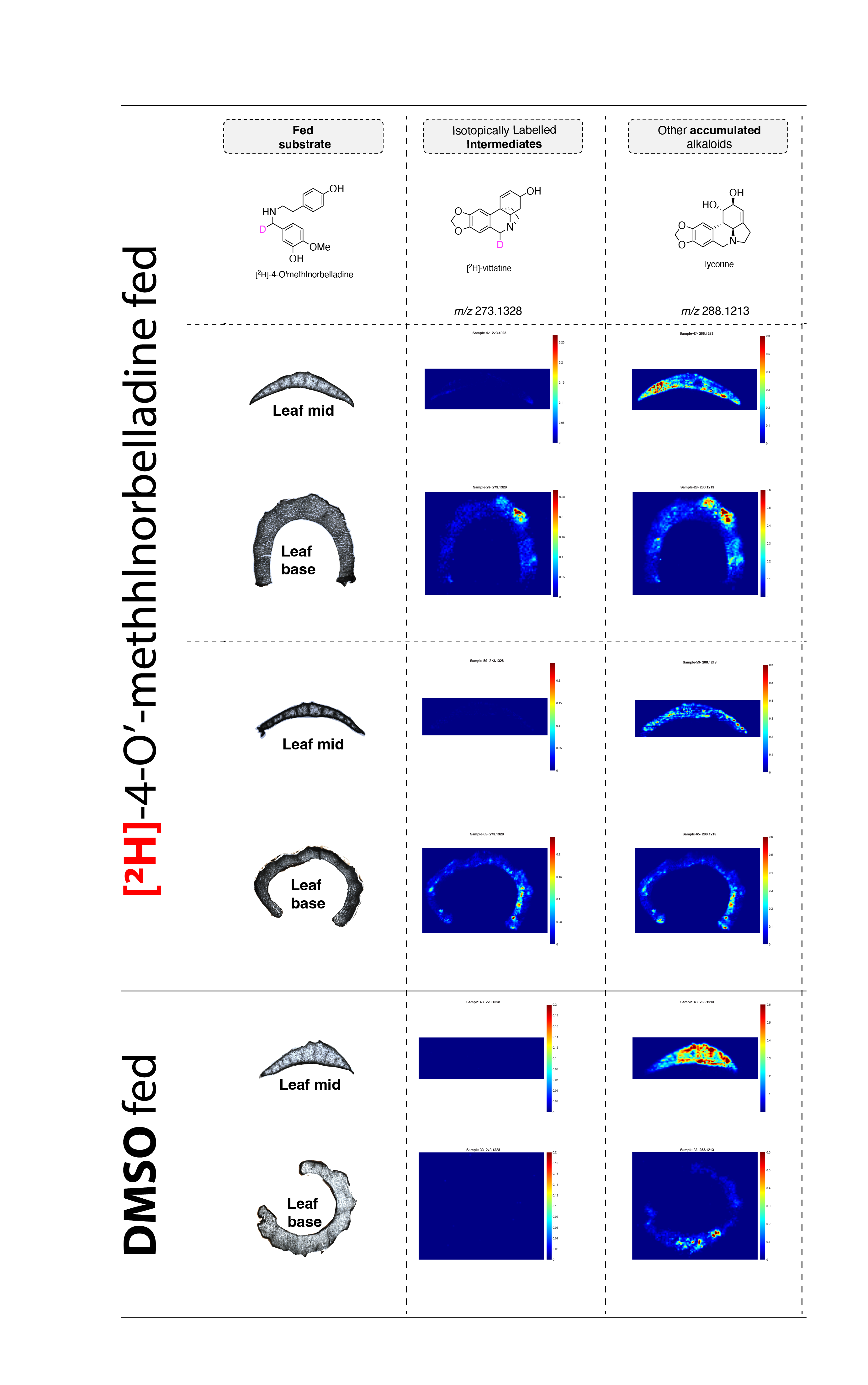

**Supplementary Figure S7:** DESI-MS images for transverse leaf sections of *Narcissus papyraceus*. Intact leaf tissues were split into two halves by cutting the leaf blade transversely (to give a top half and a bottom half). Each half was fed deuterium labelled precursor (2H-4OMN) for 7-14 days, or alternatively, a solution of 0.3% DMSO as control. The basal ~1 inch of tissue was then sectioned on a cryostat. The mass ion intensities for each sample shown here were normalized to the maximum ion intensity of lycorine within the same sample. In the labelled precursor fed sections, DESI-MS revealed that leaf bases preferentially accumulated labelled alkaloids, thus suggesting that active biosynthesis was active in leaf bases. In the DMSO-feeding control, such a trend is not observed.

**Supplementary Figure S8**: Tandem MS/MS on DESI-MS samples in Figure 2C supports the putative molecular identities of labelled vittatine and unlabelled lycorine, respectively.

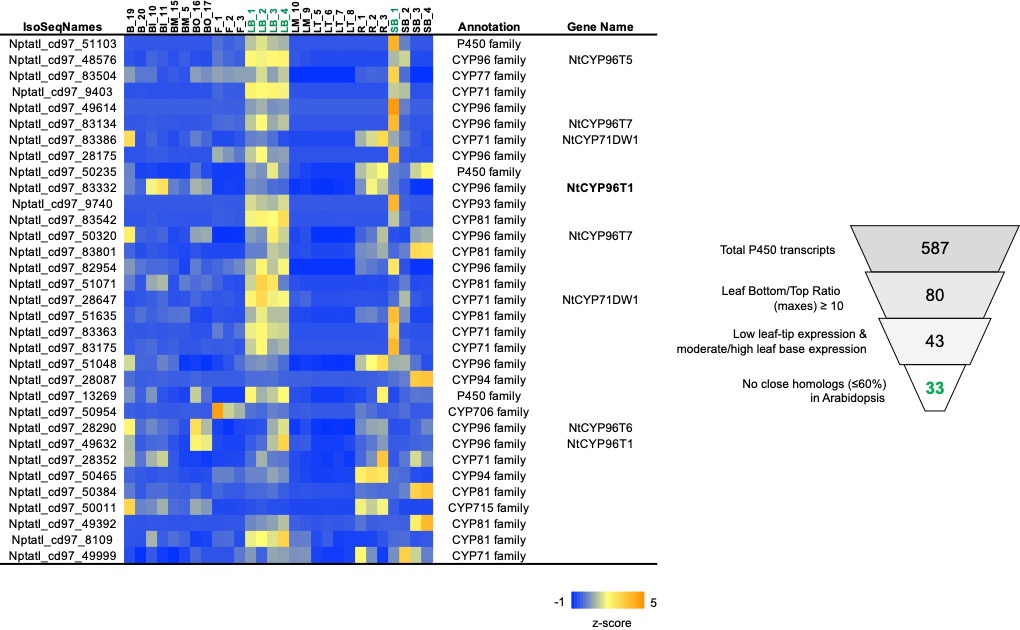

**Supplementary Figure S9:** Candidate P450s for oxidative coupling were identified by looking for cytochromes p450 that exhibited a TMM expression ratio of leaf bottom/ top of ≥ 10, low absolute leaf-top expression (≤2) and moderate/high leaf base expression (≥5), and low BLASTP %id to the closest homologs in *Arabidopsis* *thaliana* (≤60%). This resulted in the identification of 33 candidates.

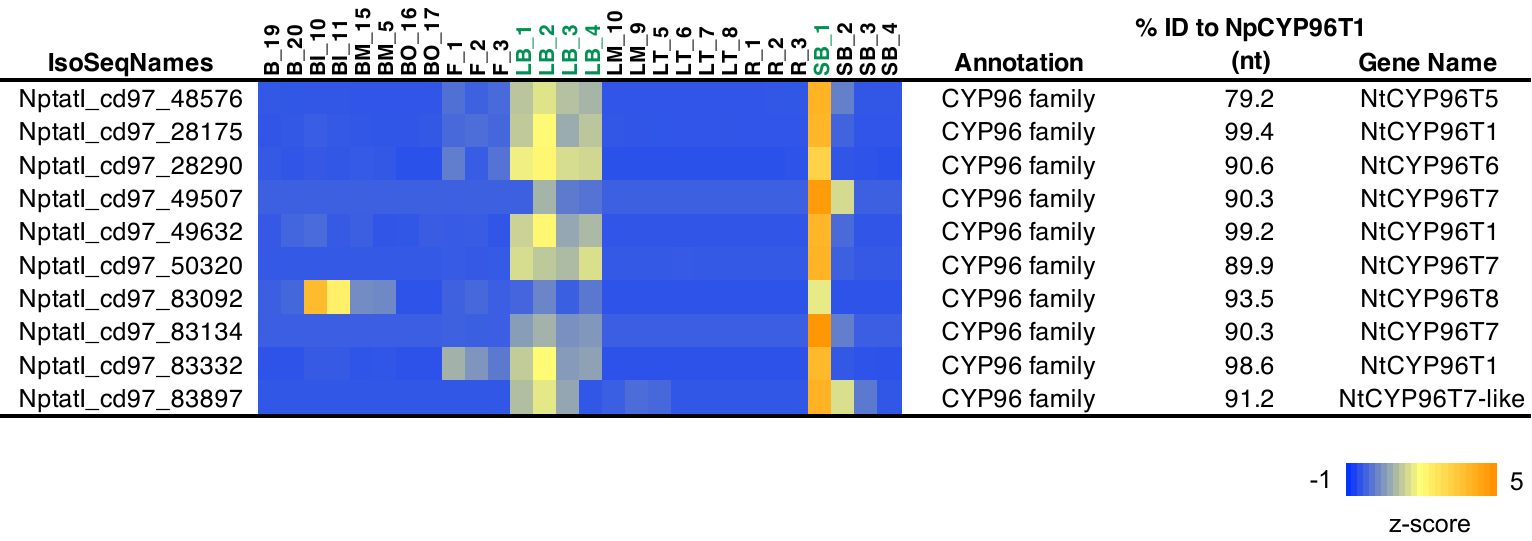

**Supplementary Figure S10:** Leaf-base expressed P450s in the long-read *Narcissus* transcriptome reported here that exhibit high homology (≥75% identity at nucleotide level) to CYP96T1

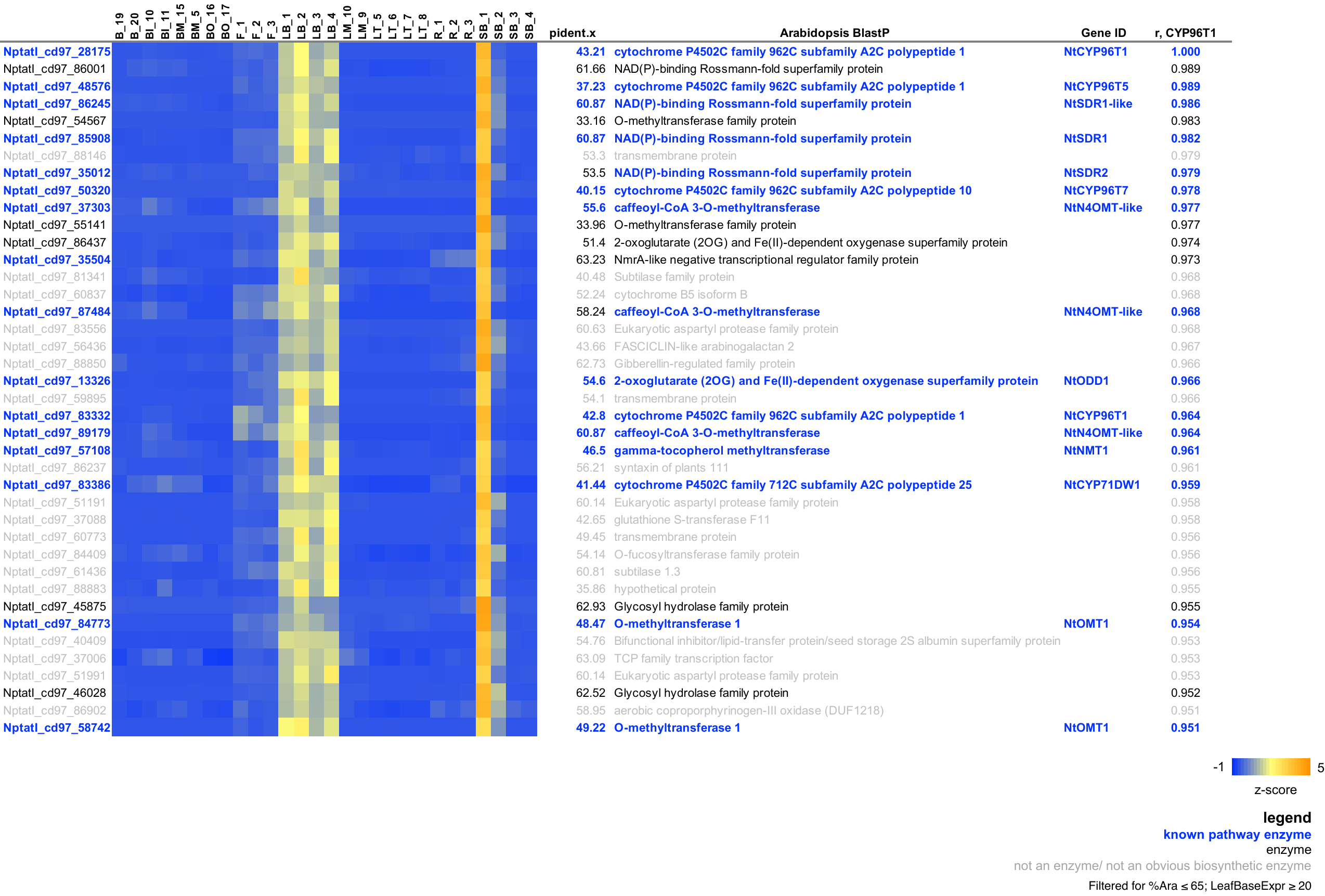

**Supplementary Figure S11**: Coexpression analysis using the p-p oxidative coupling enzyme *Np*CYP96T1 as bait reveals several candidate AmA biosynthetic enzymes. The Pearson correlation coefficient (*r*) values are listed on the right of the table. The list of coexpressing candidates was filtered for Pearson’s *r* > 0.95, blast %id to *Arabidopsis* *thaliana* homolog of ≤65%, and minimum expression at leaf bases of ≥ 20 TMM.

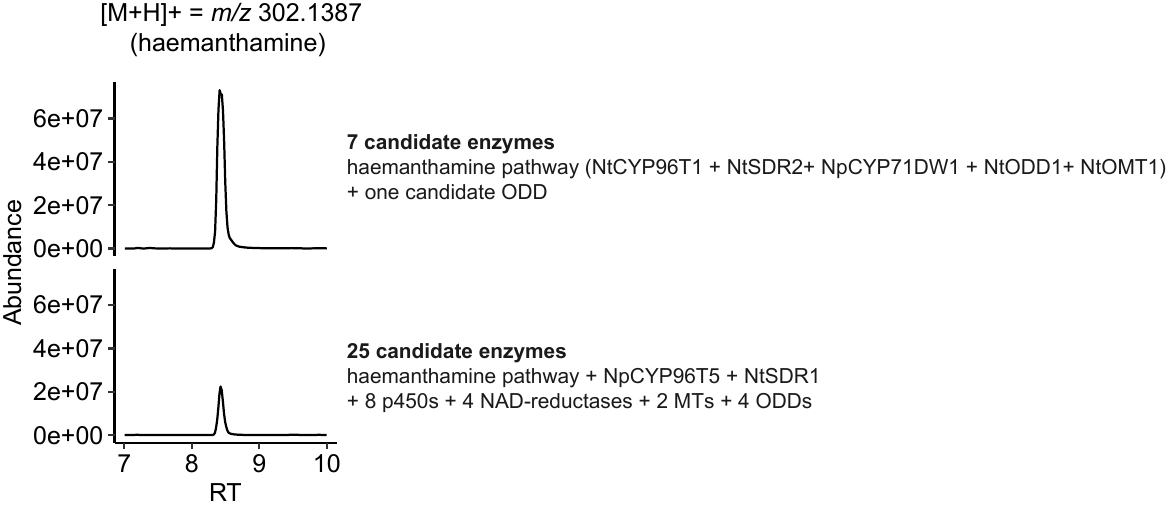

**Supplementary Figure S12:** Co-Infiltration of a total of 25 genes (i.e .the haemanthmine pathway (5 genes) alongside 17 other candidate genes) in a single leaf in our transient *N. benthamiana Agrobacterium-*mediated expression system still results in clearly detectable production of haemanthamine in the leaves.

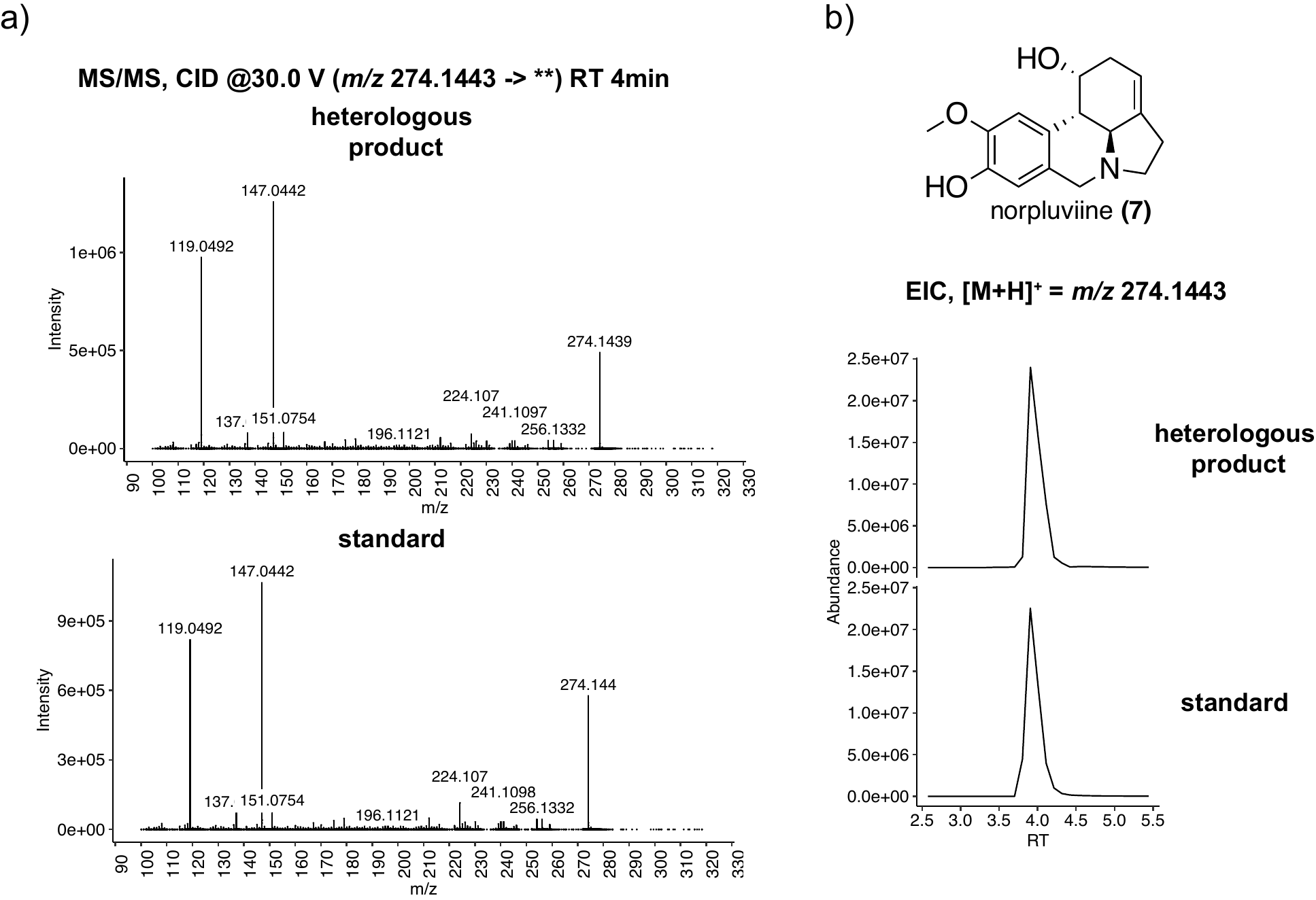

**Supplementary Figure S13:** LC-MS/MS chromatograms and MS/MS spectra of the *ortho-para’* pathway intermediate norpluviine, which was produced upon transient co-expression of *Nt*CYP96T5 and *Nt*SDR1 with 4-O’-methylnorbelladine in *N. benthamiana* as described in Methods. The identity of this mass feature was verified by comparison of retention time and MS/MS fragmentation pattern to a previously characterized authentic norpluviine standard.^1^

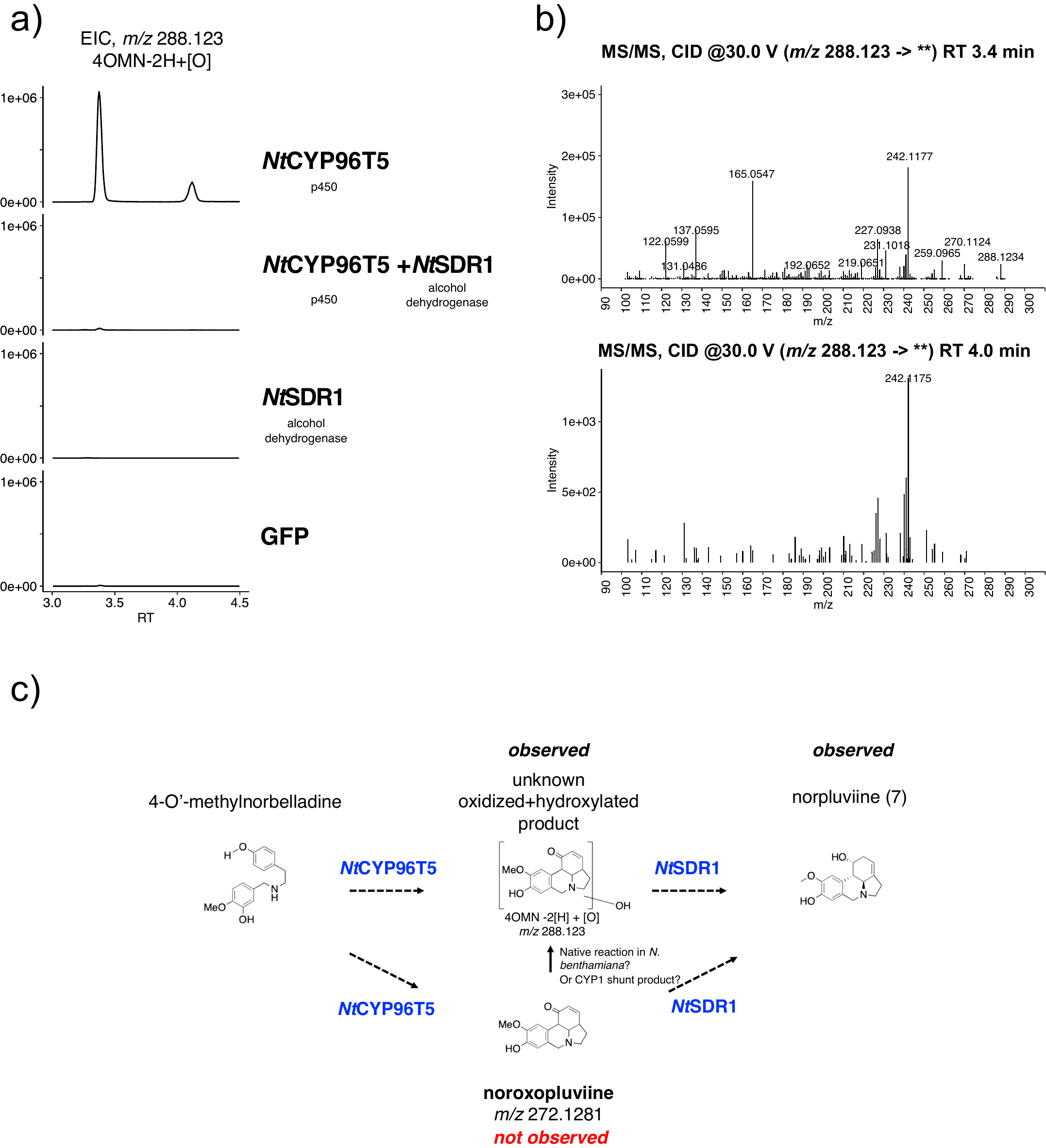

**Supplementary Figure S14:** *Nt*CYP96T5 produces an oxidized, poly-cyclic product of unknown identity that is consumed upon co-infiltration with *Nt*SDR1. (a) EICs of m/z 288.123 unknown *Nt*CYP96T5 product (b) MS/MS of the two 288.123m/z products reveals that that the product is likely poly-cyclic and possibly related to the **7** scaffold. (c) Some possible explanations for the observation of the 288.123 m/z upon infiltration of *Nt*CYP96T5 include: either, 288.123m/z is a true intermediate on pathway to **7**, or that the true product of *Nt*CYP96T5, i.e. noroxopluviine, is unstable, and is processed to the 288.123 m/z by either native enzymes present in the *N. benthamiana* expression system or some other mechanism. Alternatively, *Nt*CYP96T5 may need *Nt*SDR1 to be functional, and in the absence of *Nt*SDR1, *Nt*CYP96T5 may produce off-pathway shunt products. These data were acquired on the Agilent 6545 Q-TOF LC-MS, as described in Methods.

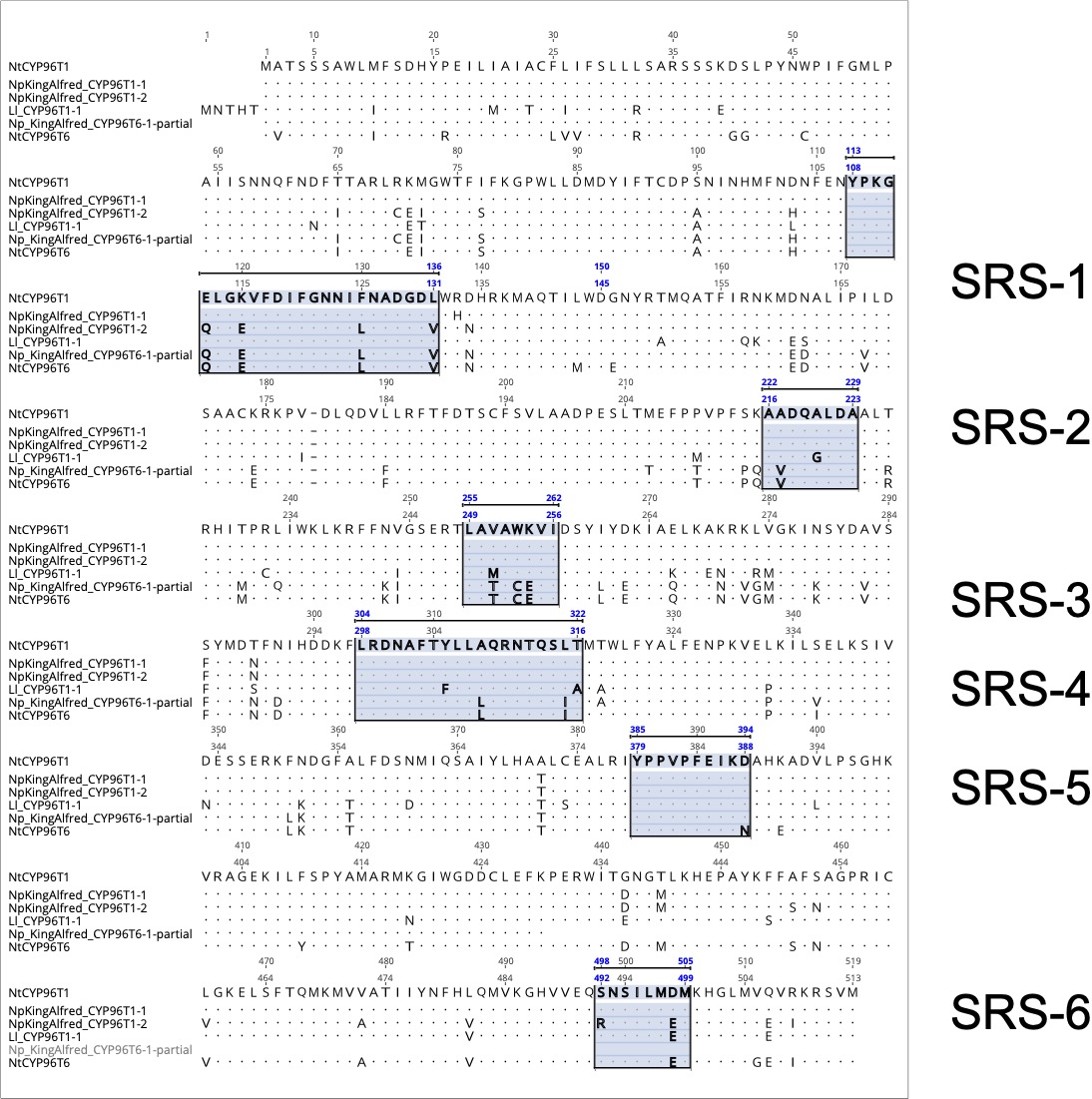

**Supplementary Figure S15:** Substrate Recognition Sites for CYP96T1 were identified through as described in Methods. The six substrate recognition sites are highlighted in blue boxes.

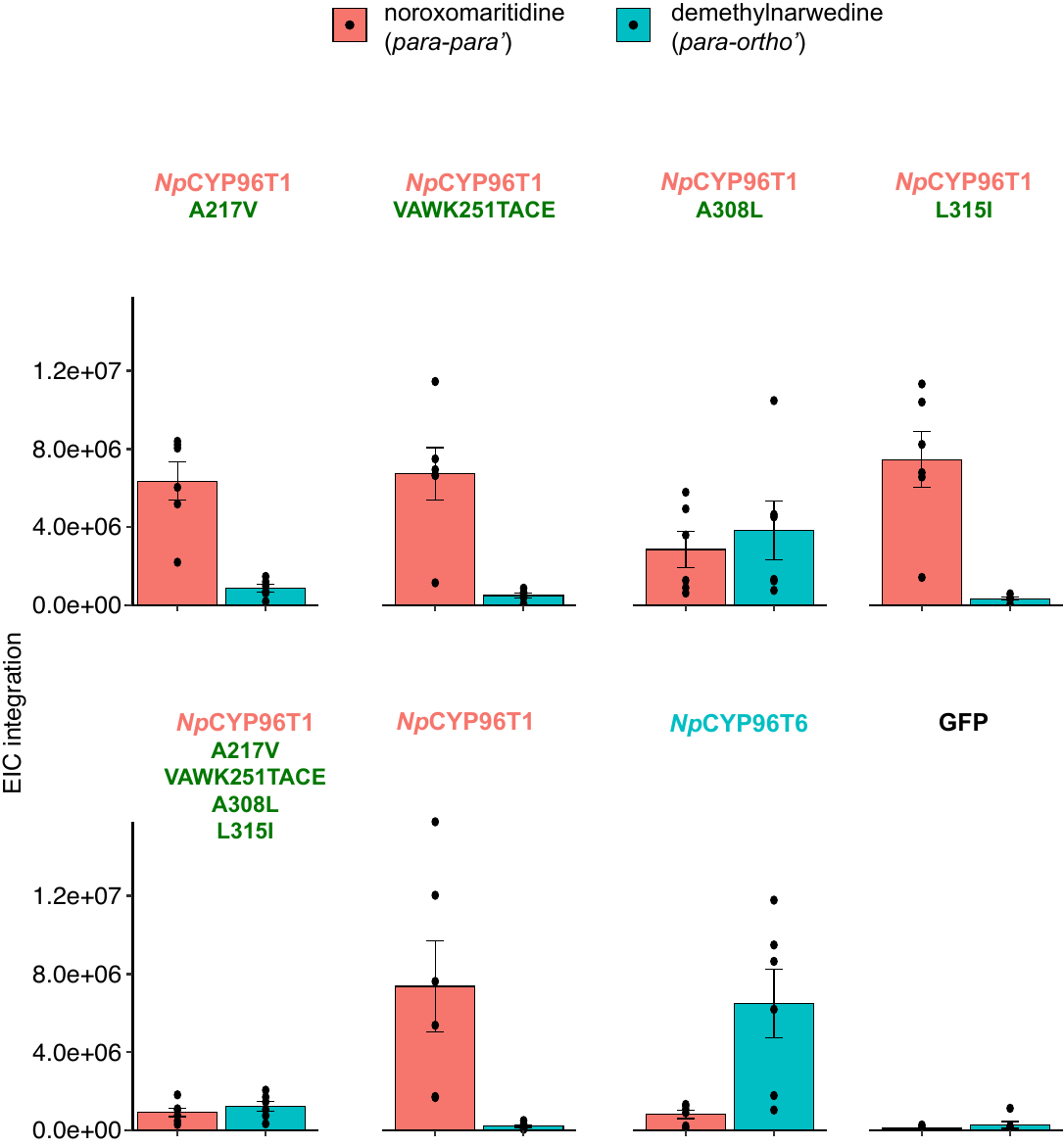

**Supplementary Figure S16:** Relative production of the *para-para’* (noroxomaritidine) and *para-ortho’* (demethylnarwedine) oxidative coupling products with infiltrated 4OMN substrate by *Nt*CYP96T1, *Nt*CYP96T6 and each of the individual mutants identified in Figure 3E (N=6 leaf replicates). These data were acquired on the Agilent 6520 Q-ToF as described in Methods.

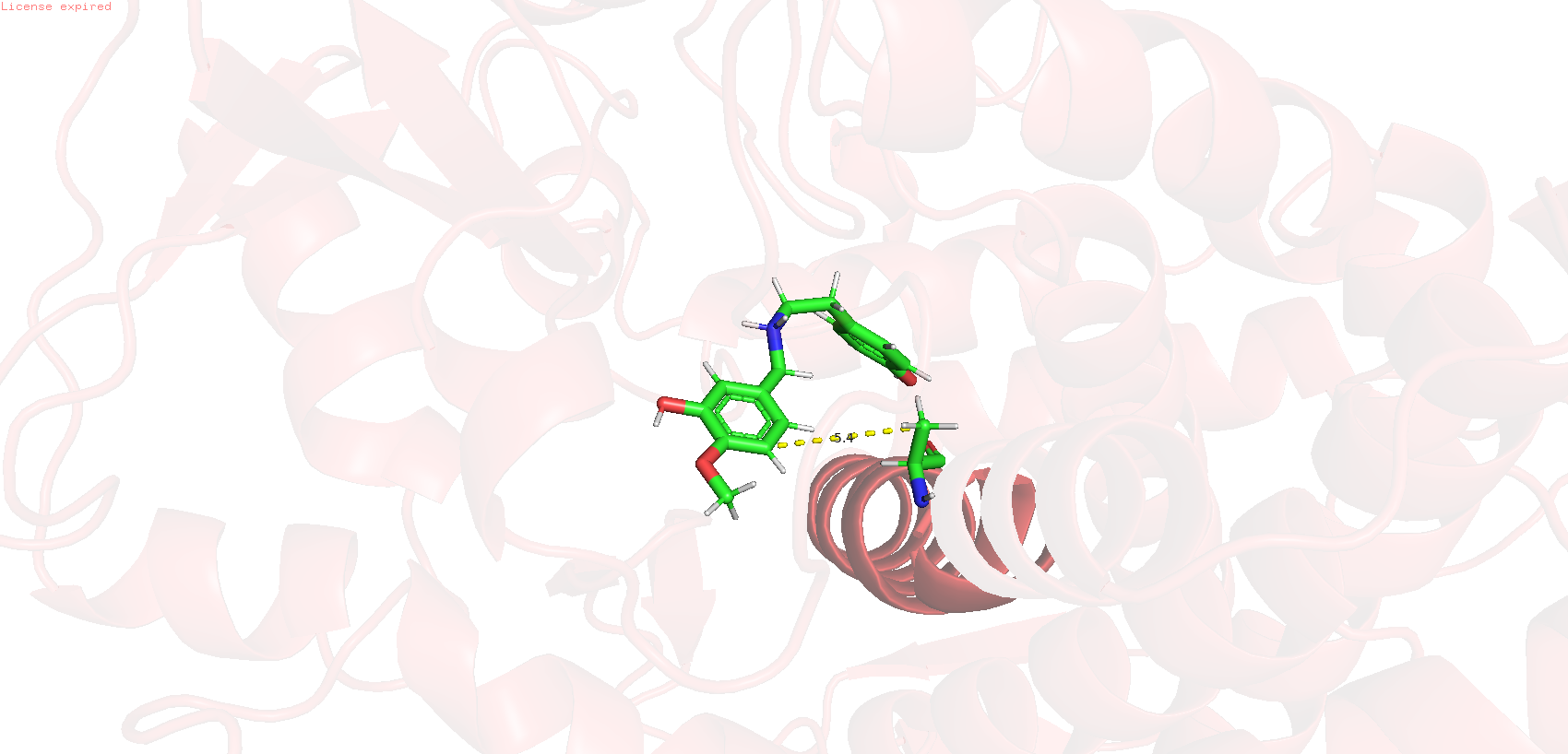

**Supplementary Figure S17:** Substrate docking of 4-O’-methylnorbelladine on a homology model of *Nt*CYP96T1 suggests that a A308, a key regioselectivity determining residue on *Nt*CYP96T1 is proximal to the 4-O’-methylnorbelladine substrate. (SWISS-model and SwissDock, against the Carotene epsilon-monooxygenase, chloroplastic, CYP97C1 template). The distance between the phenyl ring on 4-O’-methylnorbelladine and the terminal carbon on Ala308 is shown. (5.4Å).

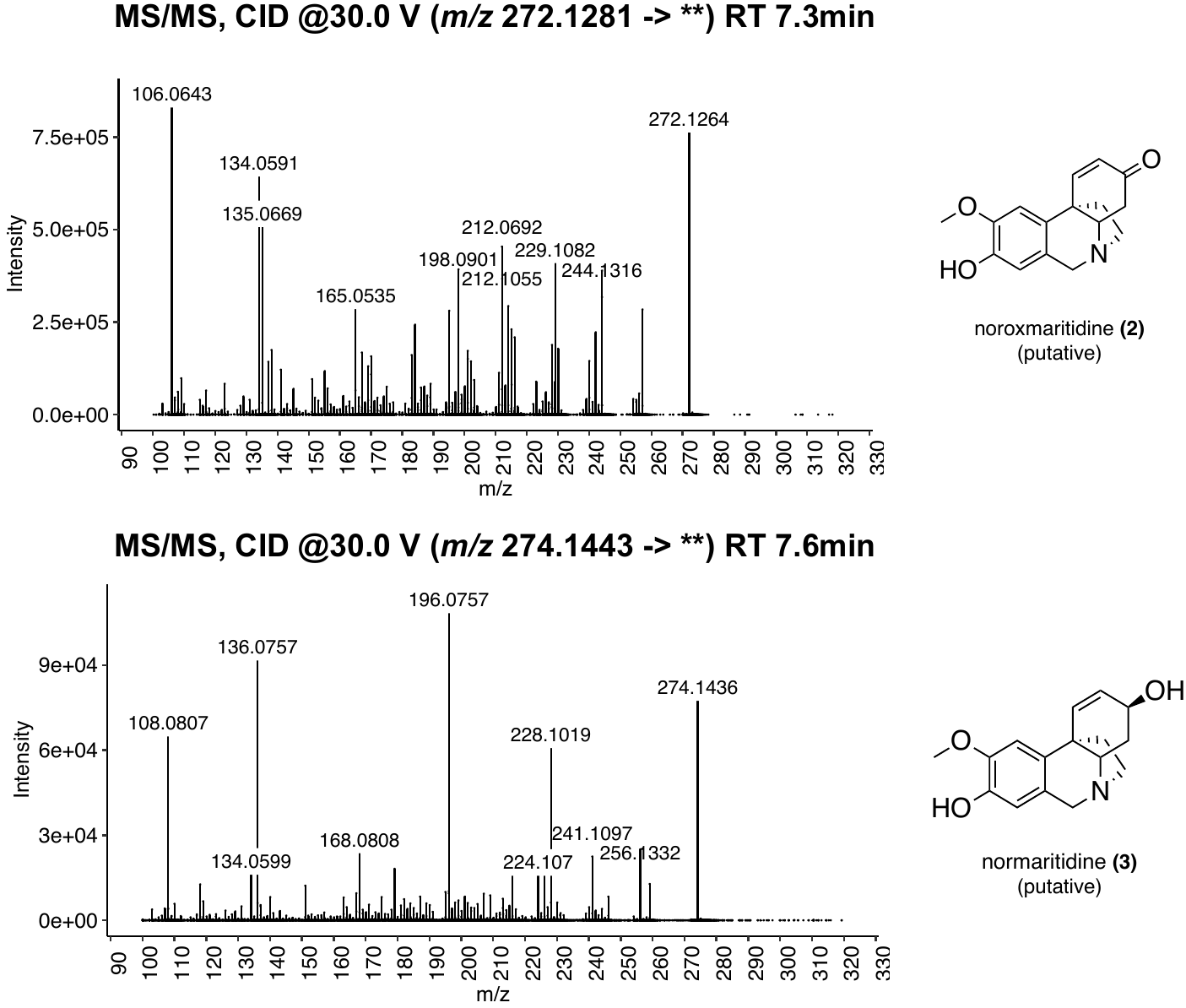

**Supplementary Figure S18:** LC-MS/MS spectra of heterologously produced (putative) *para-para’* pathway intermediates noroxomaritidine (**2**) and normaritidine (**3**). Both **2** and **3** were generated through transient co-expression of *Nt*CYP96T1 with *Nt*SDR2 with 4-O’-methylnorbelladine in *N. benthamiana* as described in Methods. These intermediates were deduced based on prior characterization of CYP96T1, expected fragmentation patterns, and expected biosynthetic intermediates on pathway to haemanthamine.^2^

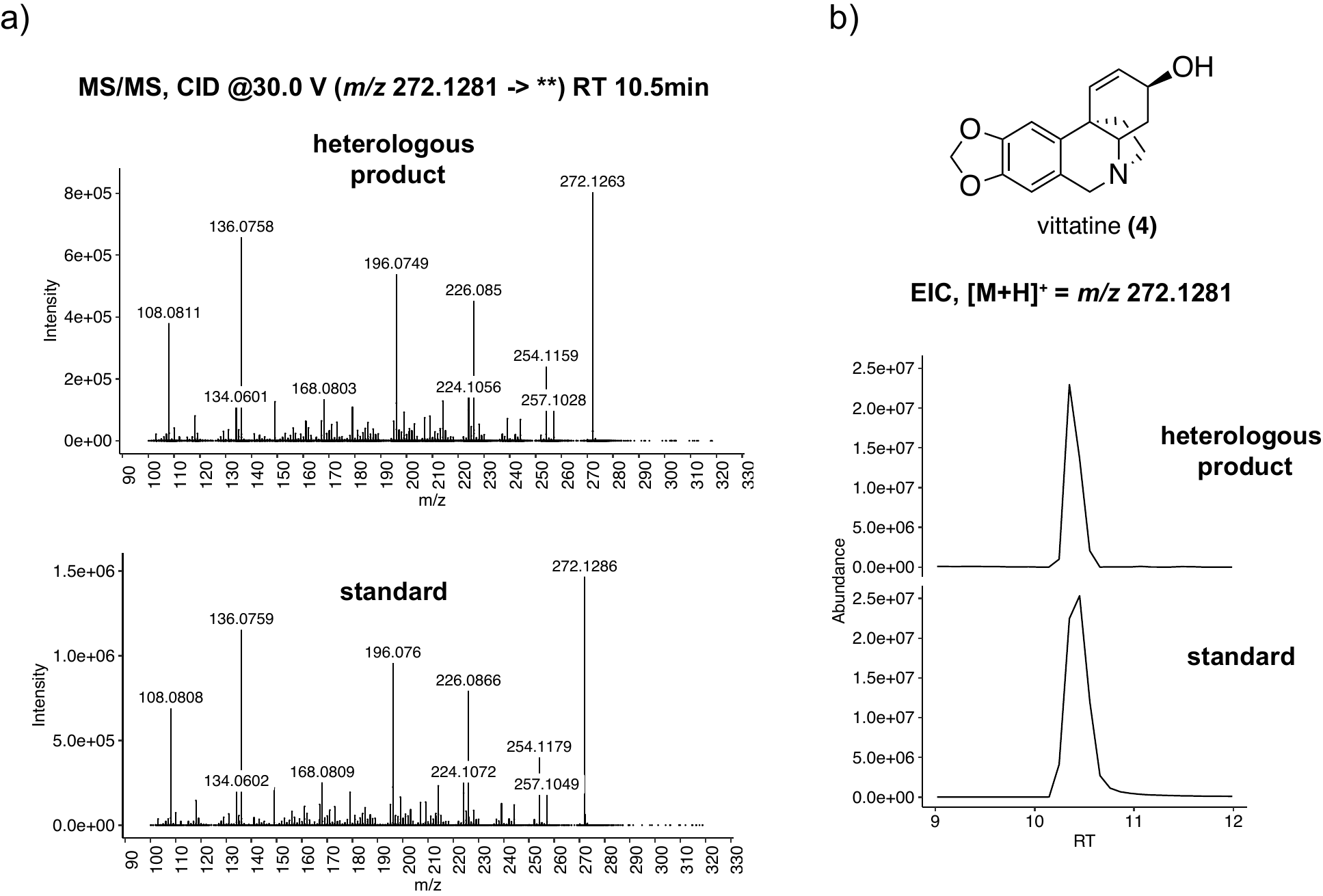
 **Supplementary Figure S19:** LC-MS/MS chromatograms and MS/MS spectra of heterologously produced *para-para’* pathway intermediate vittatine (**4**), which was observed upon transient co-expression of *Nt*CYP96T1, *Nt*SDR2 and *Nt*CYP71DW1 with 4-O’-methylnorbelladine in *N. benthamiana* as described in Methods. The identity of this mass feature was verified by comparison of retention time and MS/MS fragmentation pattern to a previously characterized authentic standard.^1^

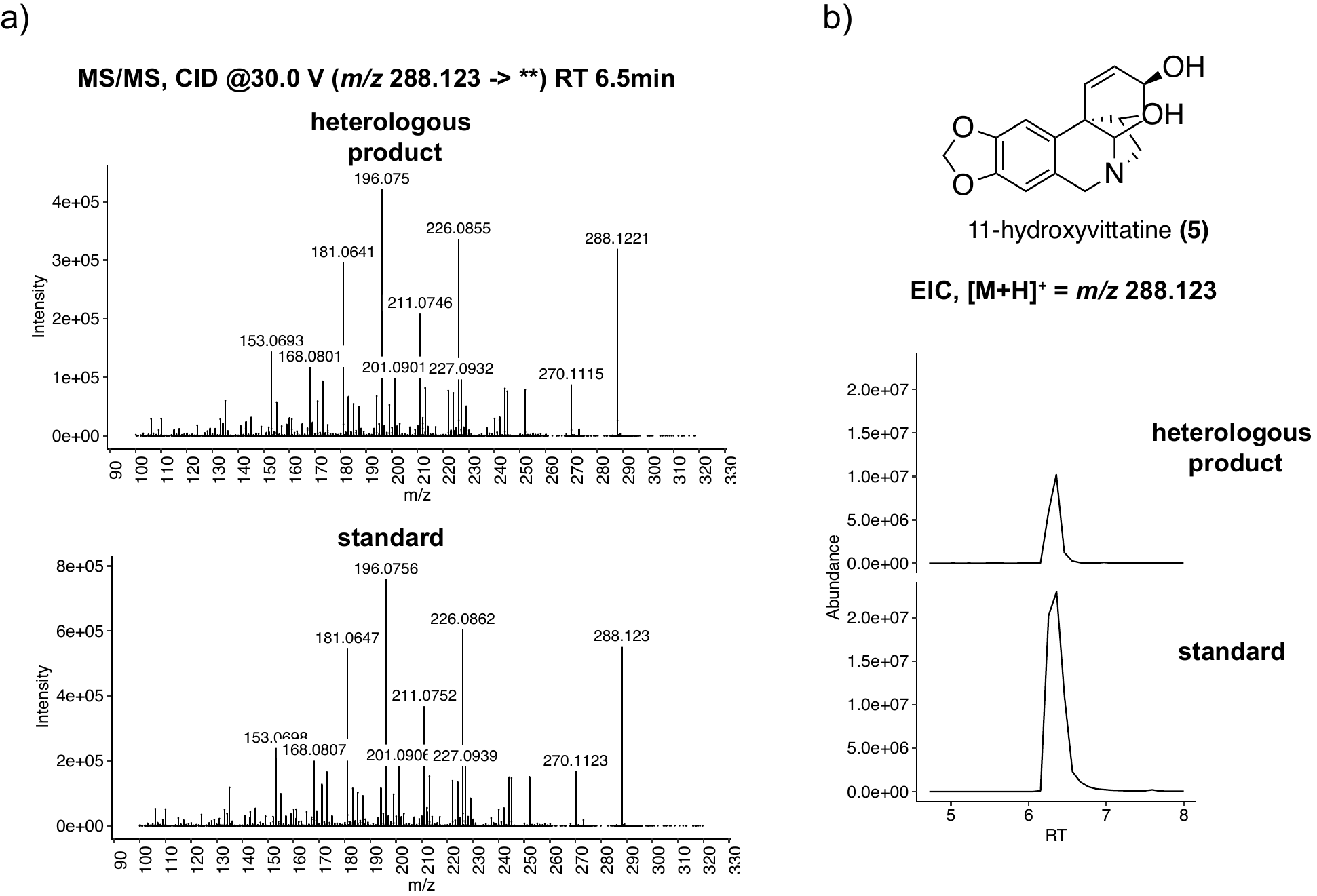

**Supplementary Figure S20:** LC-MS/MS chromatograms and MS/MS spectra of heterologously produced *para-para’* pathway intermediate 11-hydroxyvittatine (**5**), which was observed upon transient co-expression of *Nt*CYP96T1, *Nt*SDR2, *Nt*CYP71DW1 and *Nt*ODD2 with 4-O’-methylnorbelladine in *N. benthamiana* as described in Methods. The identity of this mass feature was verified by comparison of retention time and MS/MS fragmentation pattern to a previously characterized authentic **5** standard^1^.

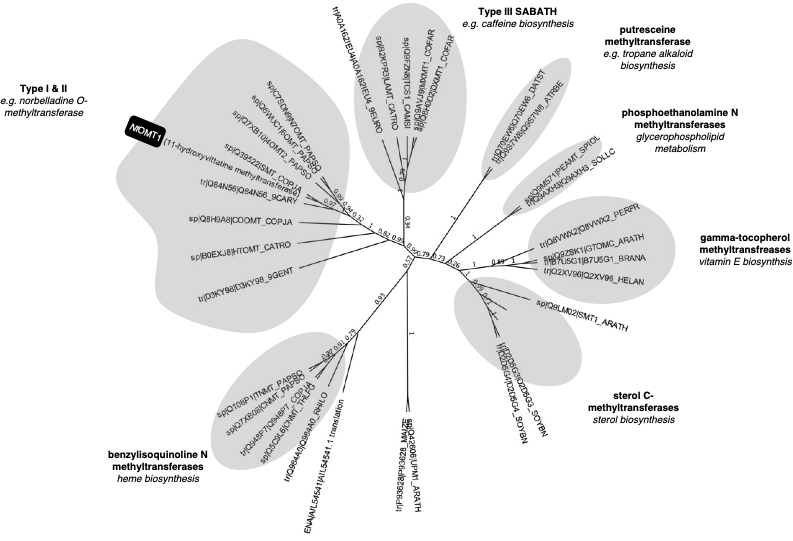

**Supplementary Figure S21:** *Nt*OMT1 belongs to the type I/II clade of plant methyltransferases. Sequences for methyltransferases were compiled from Liscombe et al, PNAS.^3^ Alignments and tree-building was performed in Geneious using clustalW and FastTree respectively. Fastree support values indicated on branches.

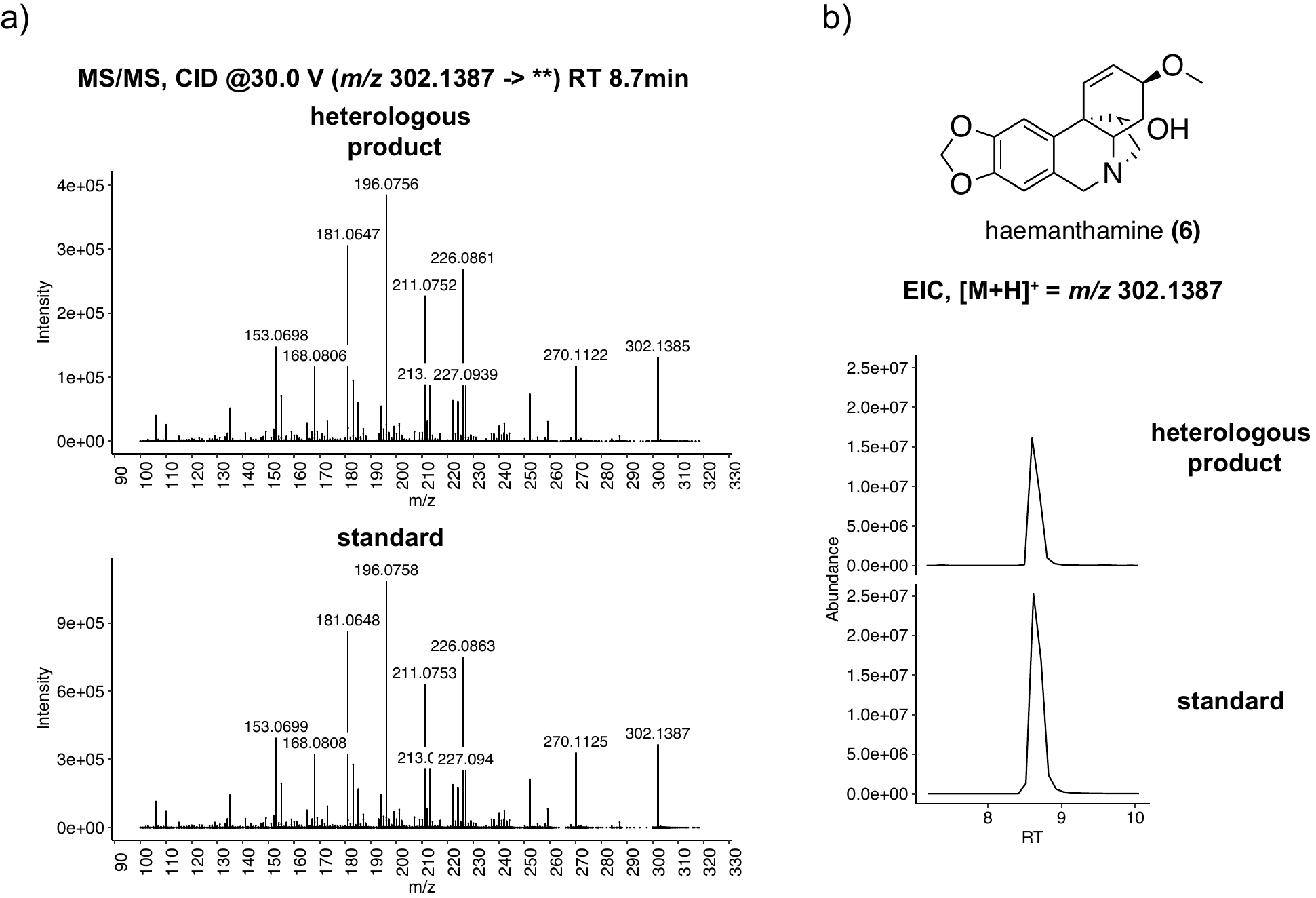

**Supplementary Figure S22:** LC-MS/MS chromatograms and MS/MS spectra of the heterologously produced eukaryotic translation inhibitor haemanthamine, which was produced upon transient co-expression of *Nt*CYP96T1, *Nt*SDR2, *Nt*CYP71DW1, *Nt*ODD2 and *Nt*OMT1 with 4-O’-methylnorbelladine in *N. benthamiana* as described in Methods. The identity of this mass feature was verified by comparison of retention time and MS/MS fragmentation pattern to a previously characterized authentic haemanthamine standard.^1^

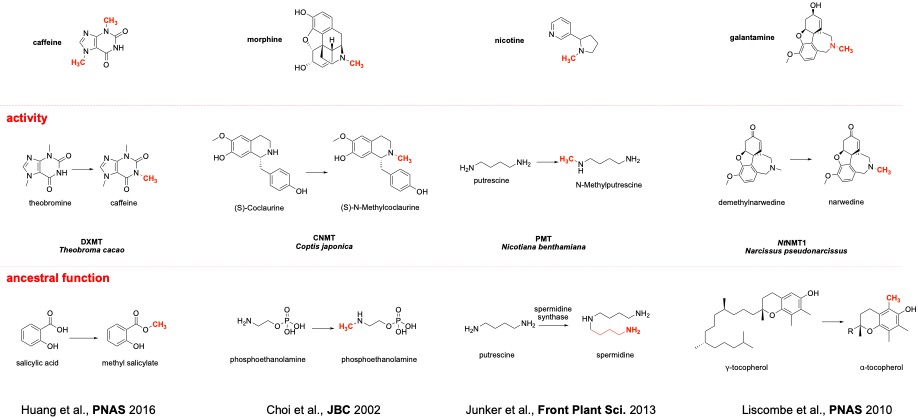

**Supplementary Figure S23**: **An overview of characterized plant N-methyltransferases** Plant N-methyltransferases are recruited from a diverse set of ancestral enzymes. Here, plant N-methyltransferases with known functions in secondary metabolism are listed, along with their presumed ‘ancesteral’ function from which they were specialized.^3–7^

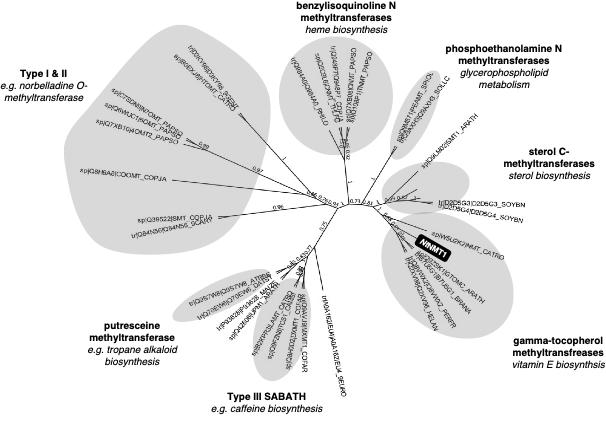

**Supplementary Figure S24**: NtNMT1 is a γ-tocopherol methyltransferase. Sequences for methyltransferases were complied from Liscombe PNAS.^8^ Alignments and tree-building was performed in Geneious using clustalW and FastTree respectively. Fastree support values indicated on branches.

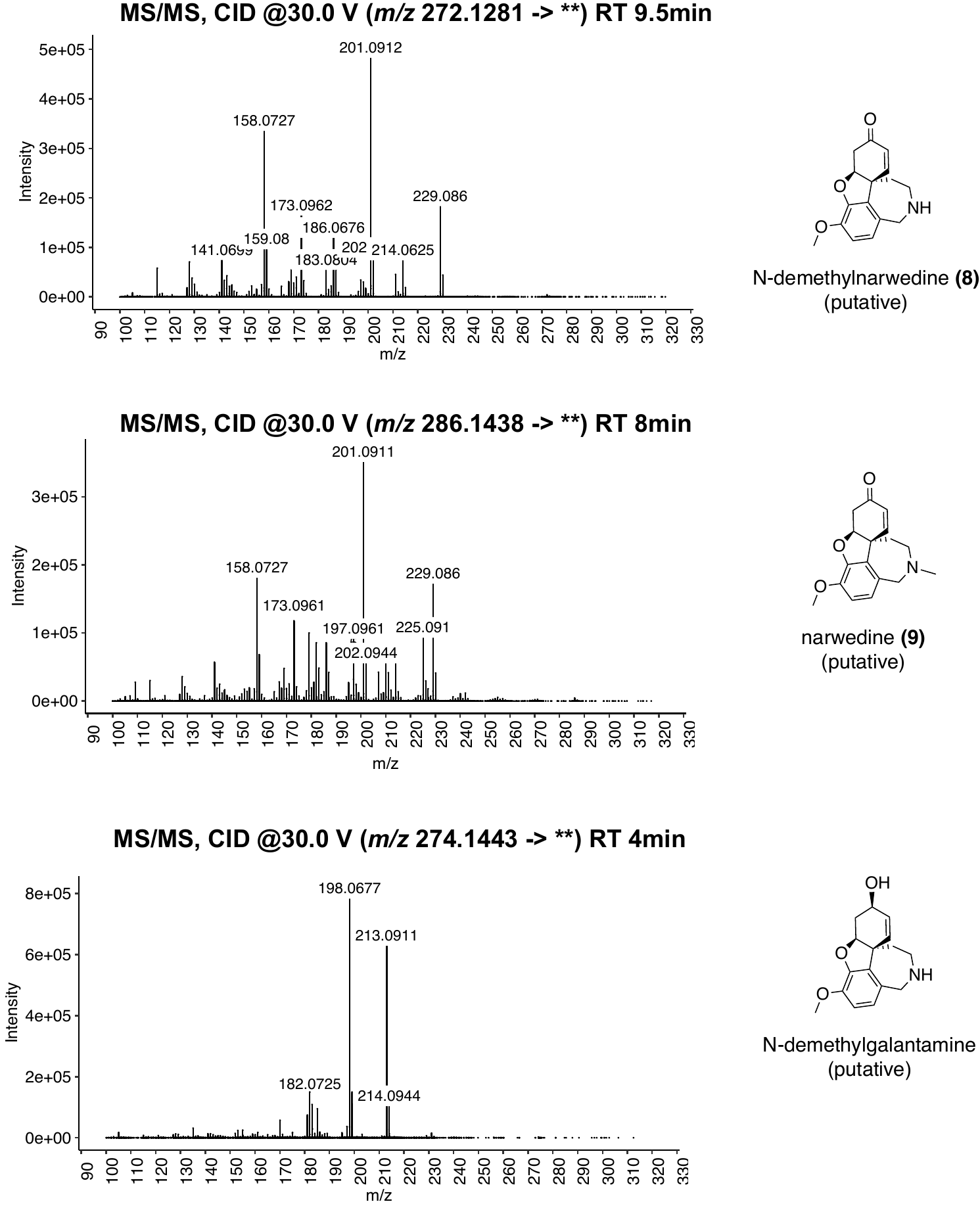

**Supplementary Figure S25:** LC-MS/MS spectra of heterologously produced (putative) *para-para’* pathway intermediates N-demethylnarwedine, narwedine, and N-demethylgalantamine. The putative N-demethylnarwedine product was generated through transient agroinfiltration of *Nt*CYP96T6 with 4-O’-methylnorbelladine in *N. benthamiana* as described in Methods, whereas the putative narwedine was observed upon transient co-expression of *Nt*CYP96T6 and *Nt*NMT1. The production of demethylgalantamine was observed upon co-infiltration of *Nt*CYP96T6 and *Nt*AKR1, as further elaborated in Supplementary Figure S27. These intermediates were deduced based on expected fragmentation patterns, prior literature reports and expected biosynthetic intermediates on pathway to galantamine.^9^

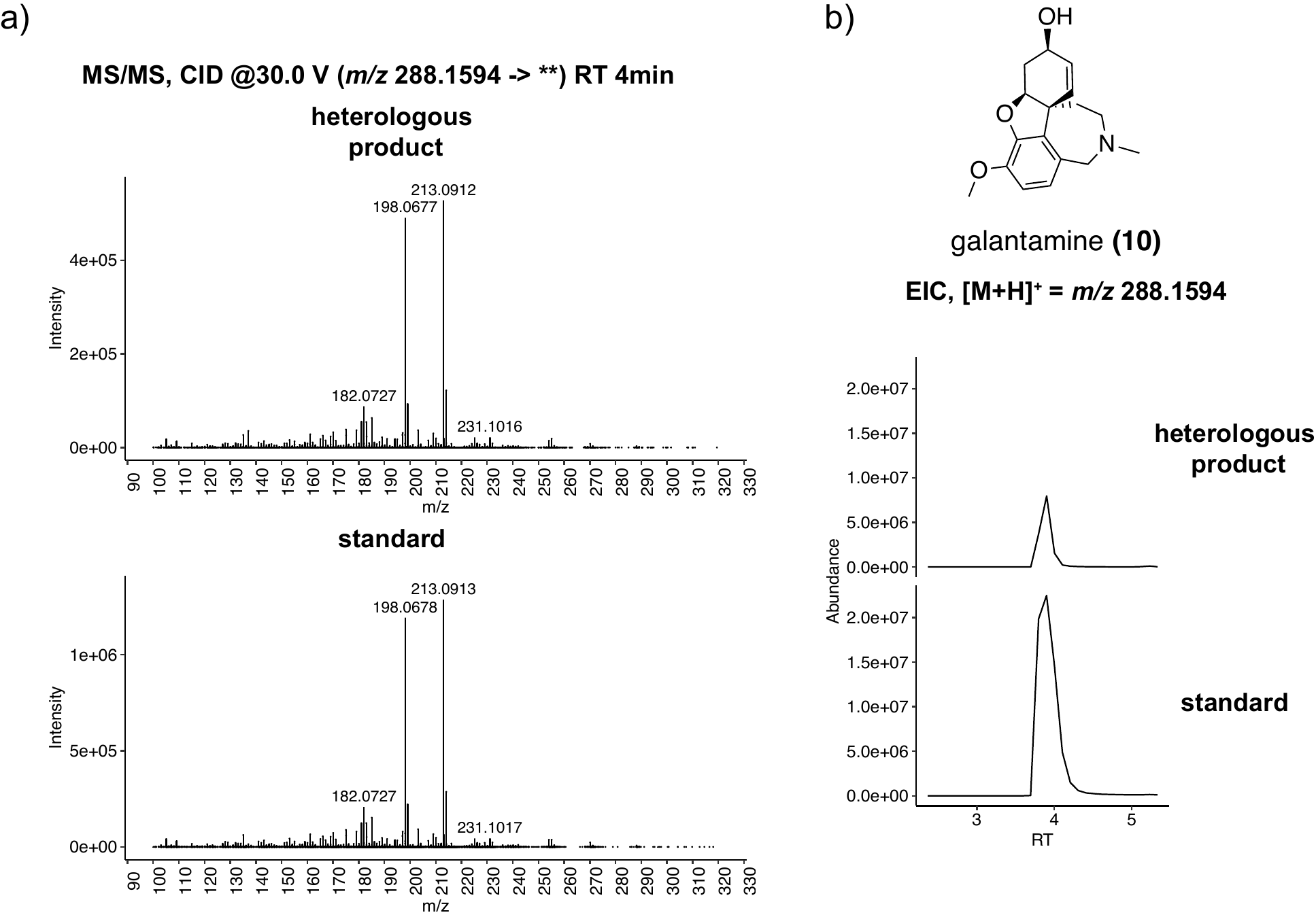

**Supplementary Figure S26:** LC-MS/MS chromatograms and MS/MS spectra of the heterologously produced galantamine, production of which was observed upon transient co-expression of *Nt*CYP96T6, *Nt*NMT1, and *Nt*AKR1 with 4-O’-methylnorbelladine in *N. benthamiana* as described in Methods. The identity of this mass feature was verified by comparison of retention time and MS/MS fragmentation pattern to a previously characterized authentic haemanthamine standard.^1^

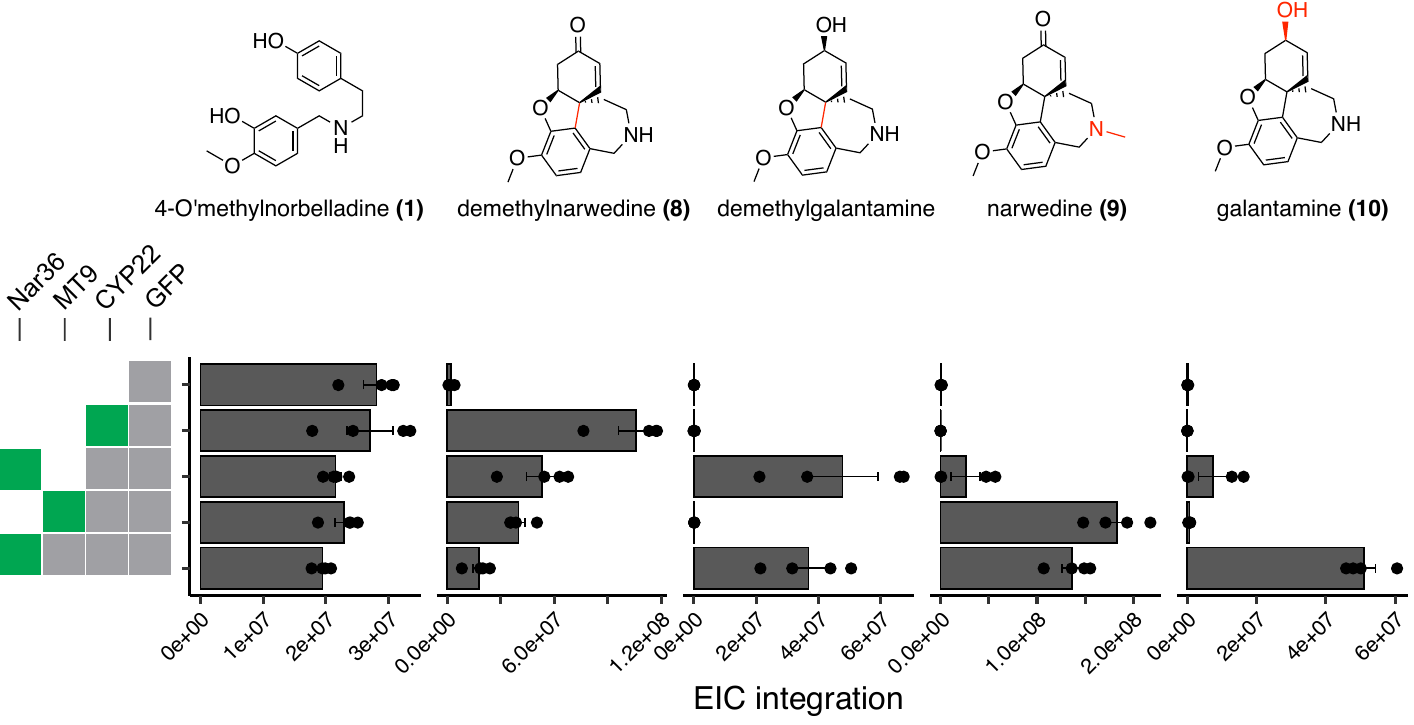

**Supplementary Figure S27:** *Nt*AKR1 can process the *Nt*CYP96T6 product demethylnarwedine (**8**) to produce demethylgalantamine. The higher observed consumption of **8** after the addition of both *Nt*NMT1 and *Nt*AKR1 (as opposed to just *Nt*NMT1 or *Nt*AKR1) may suggest that galantamine biosynthesis may involve both demethylgalantamine and narwedine (**9)** as intermediates. The EIC integrations were calculated for [M+H]^+^ ions for each of the displayed pathway intermediates.

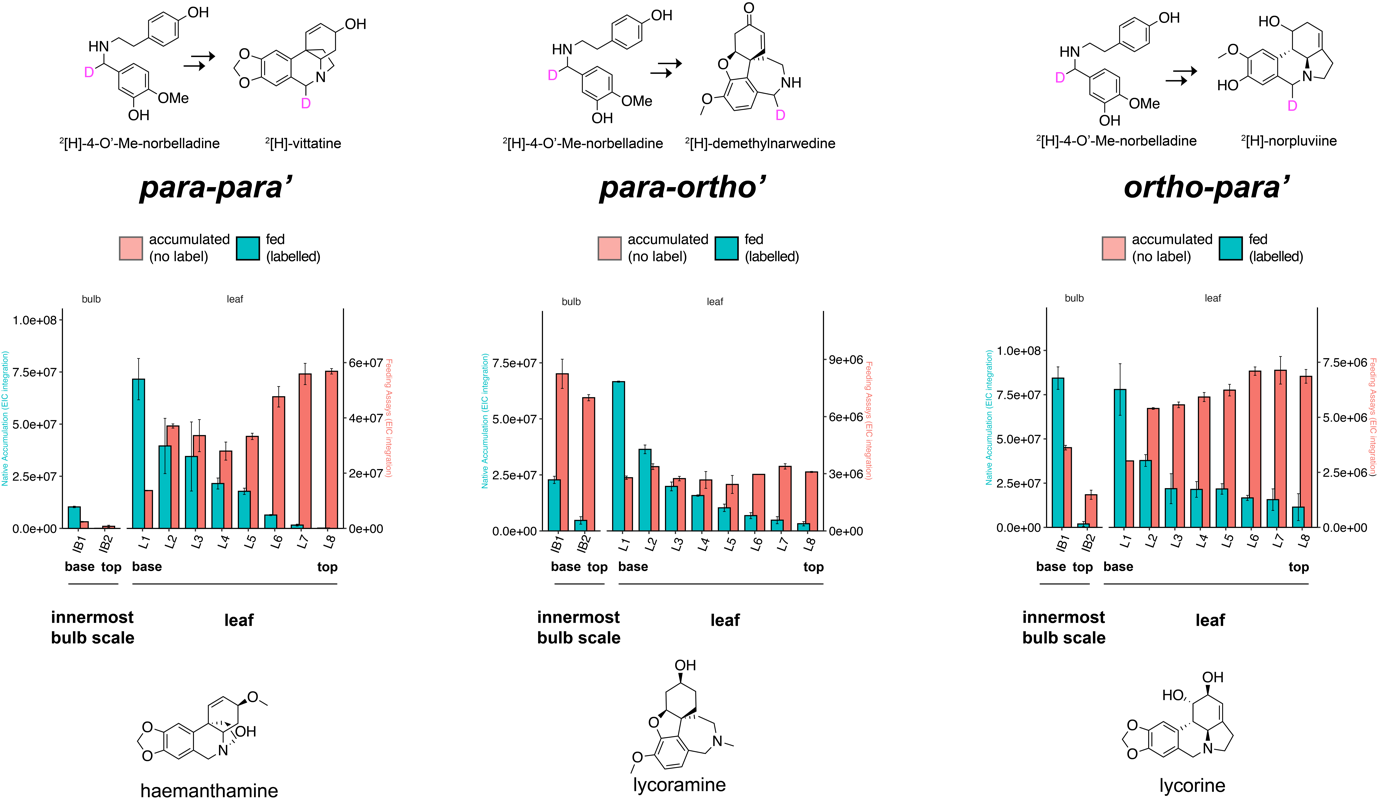

**Supplementary Figure S28**: All three branches of the AmA pathways are active in leaf and bulb bases. This figure is an extension of Figure 2, with feeding and accumulation levels for innermost bulb scales included. The extracted ion abundances (peak integration) are shown for: unlabelled haemanthamine (*m/z* 303.1426 [(M+1)+H]^+^), unlabelled lycoramine (*m/z* 291.1790 [(M+1)+H]^+^), unlabelled lycorine (*m/z* 289.1270 [(M+1)+H]^+^), deuterium labelled ^2^[H]-norpluviine (*m/z* 275.1506, [M+H]^+^), deuterium labelled ^2^[H]-demethylnarwedine(*m/z* 273.1349, [M+H]^+^), deuterium labelled ^2^[H]-vittatine (*m/z* 273.1349, [M+H]^+^). N=2 biological replicates were used for all feeding and metabolite quantification experiments. All experiments measuring the accumulation of un-labelled intermediates here were run on separate sets of leaves from the same plant and were not fed any precursors. The M+1 isotopologue for the natively accumulated, unlabelled alkaloids haemanthamine, lycoramine and lycorine were quantified to avoid detector saturation effects, as these alkaloids were present in very high abundances in our assays.

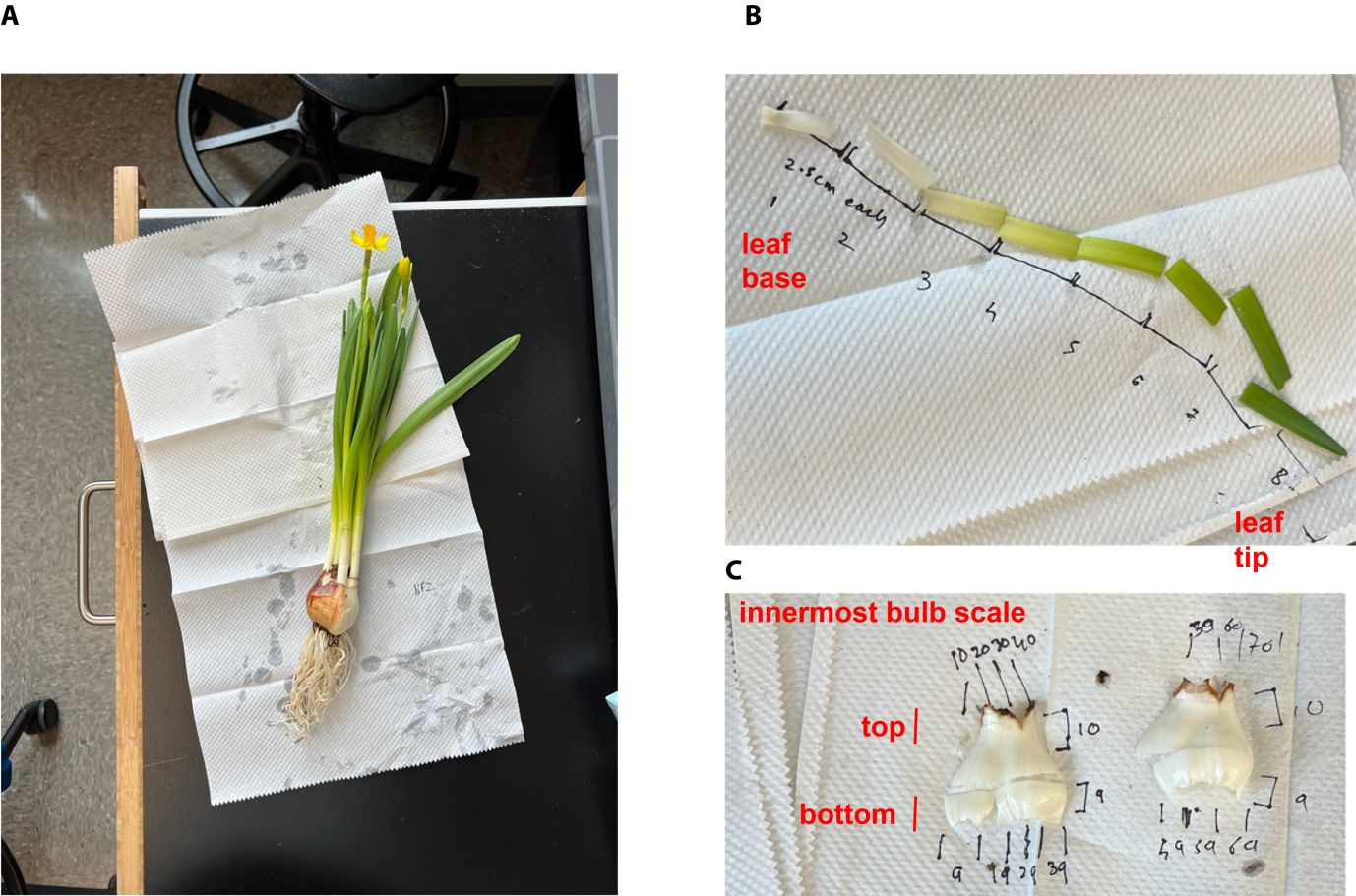

**Supplementary Figure S29**: Harvesting leaf and bulb tissue sections for experiments reported in Figure 2. (A) A fully mature *N. pseudonarcissus tête-à-tête* daffodil was used for feeding, metabolomics and RNA-extarctions. (B) Eight sections of approximately 2.5cm each from leaves of a single fully mature daffodil were used for metabolomics, feeding and RNA extraction. (C) The innermost bulb scale that was not the base of a current foliage leaf was also divided into two, and similarly processed as leaves for metabolomics analysis, RNA extraction and feeding studies. The leaf tip (8) represents the oldest part of the leaf, and the leaf base (1) represents the youngest. Similarly, the bottom of the bulb scale is the younger section of the scale.

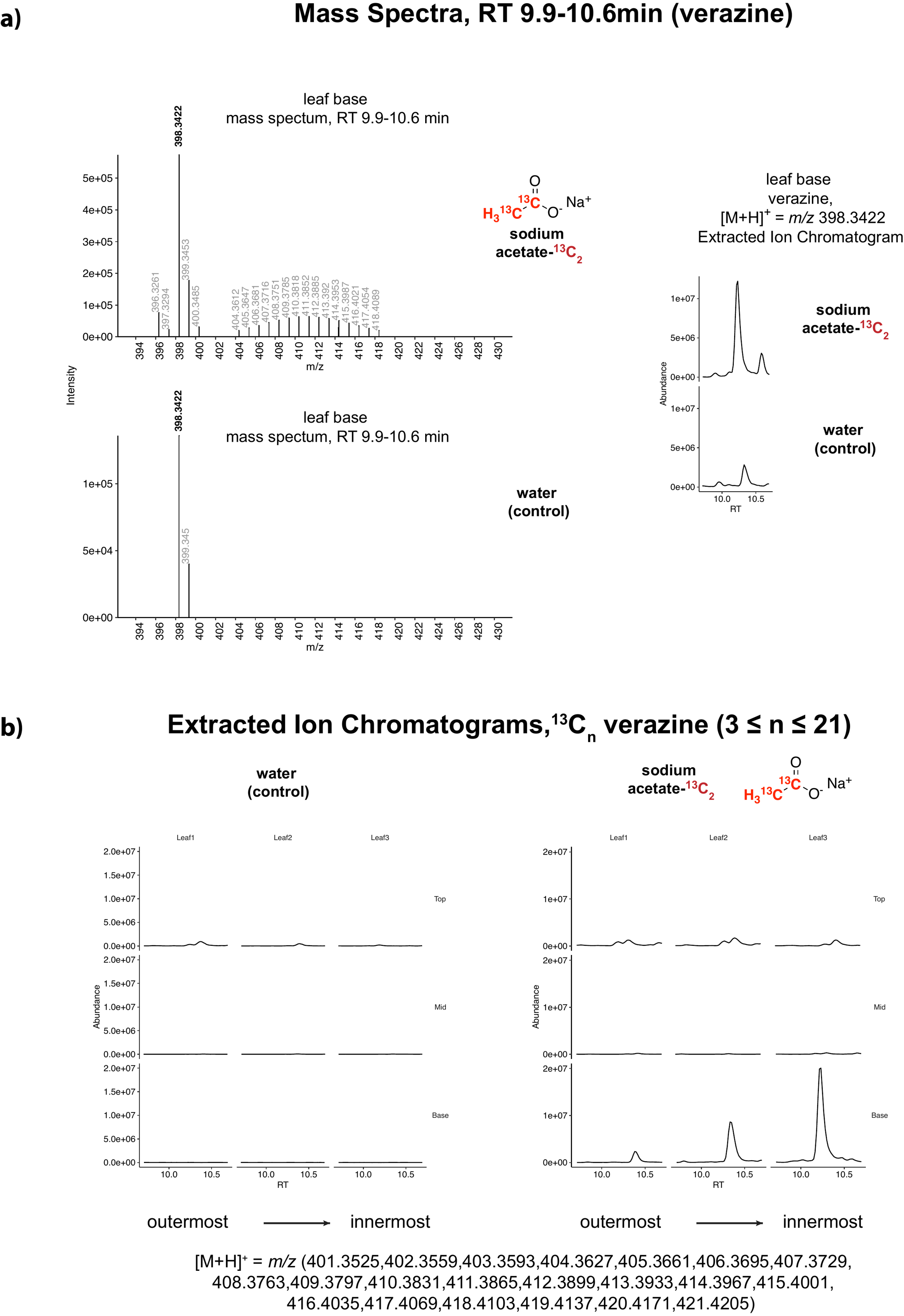

**Supplementary Figure S30: A)** Feeding studies on *Veratrum* *nigrum* plants with labelled sodium acetate-^13^C_2_ reveal incorporation of isotope label into a mass feature that matches the MS/MS fragmentation pattern for verazine as reported previously in literature (Supplementary Figure S31). The isotope incorporation presents itself as an envelope of isotopic masses corresponding to different numbers of ^13^C units incorporated into the putative verazine mass feature. **B)** The level of incorporation of the isotope label into verazine increases from the outermost leaf to the innermost leaf, which may reflect the differences in age and development stage for each of the leaves. A combined EIC is reported here for all the observed masses in the labelled ^13^C_n_-verazine (3 ≤ n ≤ 21) isotopic envelope.

**Supplementary Figure S31:** MS/MS supports the isotopic labelling of verazine. The MS/MS fragmentation patterns for the parent unlabelled mass (top row), [M+H]^+^ *= m/z* 393.3423 is consistent with previous reports of the MS/MS fragmentation patterns for verazine, with *m/z* 126.1277 and *m/z* 159.1169 as major fragments^10^. Moreover, the fragmentation patterns of other mass ions within the (labelled) isotopic envelope of verazine are consistent with the expected fragmentation patterns for isotopically labelled verazine, thus suggesting that the isotopic envelope observed for verazine in Supplementary Figure S30A indeed corresponds to ^13^C-labelled verazine.

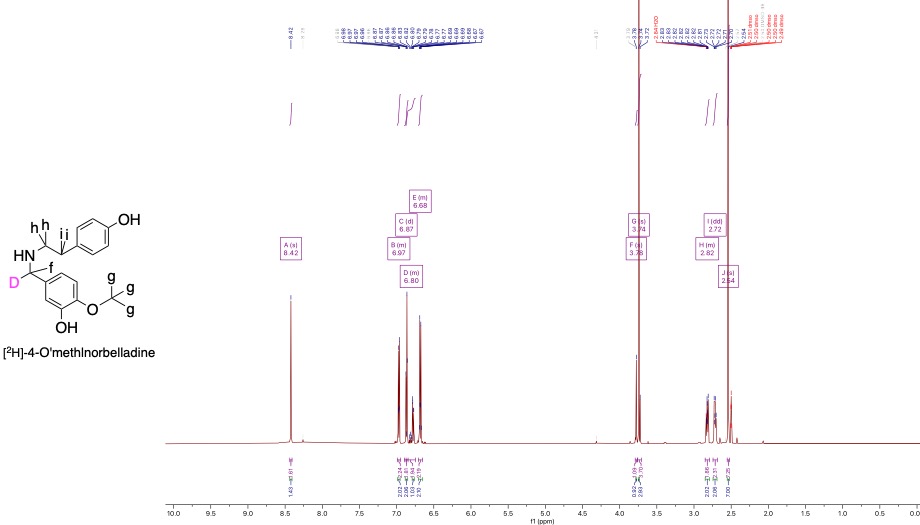

**Supplementary Figure S32:** ^1^H-NMR spectrum of synthetic [^2^H]-4-O’-methylnorbelladine used for feeding studies. [^2^H]-4-O’-methylnorbelladine: ^1^H NMR (600 MHz, DMSO-d_6_) δ 8.42 (s, 2H), 6.99-6.95 (m, 2H), 6.87 (d, J = 8.19 Hz, 2H), 6.88-6.75 (m, 2H), 6.71-6.65(m, 2H), 3.78(s, 1H), 3.74(s, 4H), 2.85-2.79(m, 2H), 2.72(dd, J = 6.45, 9.6 Hz, 2H). The sample contained a small amount of DMSO carried over from synthesis (Methods) that appears as a single peak (peak J). Putative proton assignments for peaks have been indicated in the chemical structure on the left, where each lower-case alphabet corresponds to a proton for the corresponding upper-case alphabet peak on the spectrum. Shifts are referenced to the residual solvent peak (DMSO, Acros Organics). Multiplicities are described as, s = singlet, d = doublet, dd = doublet of doublets, dt = doublet of triplets, m = multiplet.

| **Sample Name** | **Tissue Section** | **Illumina sequencing?** | **PacBio sequencing?** |
| --- | --- | --- | --- |
| F-1 | inflorerscence 1 | y |  |
| F-2 | inflorerscence 2 | y | Y |
| F-3 | inflorerscence 3 | y |  |
| SB-1 | stem bottommost | y | y |
| SB-2 | stem lower mid | y |  |
| SB-3 | stem upper mid | y |  |
| SB-4 | stem top | y |  |
| R-1 | roots 1 | y |  |
| R-2 | roots 2 | y |  |
| R-3 | roots 3 | y |  |
| LT-5 | leaf 1, top | y |  |
| LT-6 | leaf 2, top | y | y |
| LT-7 | leaf 3, top | y |  |
| LT-8 | leaf 4, top | y |  |
| LB-1 | leaf 1, bottom | y |  |
| LB-2 | leaf 2, bottom | y | y |
| LB-3 | leaf 3, bottom | y |  |
| LB-4 | leaf 4, bottom | y |  |
| LM-1 | leaf 1, mid | y |  |
| LM-2 | leaf 2, mid | y |  |
| BI-10 | bulb scale, innermost | y |  |
| BI-11 | bulb scale, innermost | y |  |
| BI-12 | bulb scale, innermost | y |  |
| BM-13 | bulb scale, mid-inner | y |  |
| BM-14 | bulb scale, mid-inner | y |  |
| BM-5 | bulb scale, mid-inner | y |  |
| BO-16 | bulb scale, outermost | y | y |
| BO-17 | bulb scale, outermost | y |  |
| BO-18 | bulb scale, outermost | y |  |
| B-19 | bulb scale, mid-outer | y |  |
| B-20 | bulb scale, mid-outer | y |  |

**Supplementary Table 1:** Sample abbreviations and descriptions for RNA-seq of *N. pseudonarcissus* tissue. See methods section for description of plant tissue, RNA extraction and sequencing protocols used. Samples that were additionally used for PacBio IsoSeq are indicated.

| **Mutant Name** | **Primer1** | **Primer2** | **Template Used** |
| --- | --- | --- | --- |
| Nar_161 | TCTGCCCAAATTCGCGACCGGTATGGTCACTTCTTCTTCAGC | TTGAGCTTCCAAATCAGGC | Nar_160 |
|  | CCGCGCCTGATTTGGAA | GAAACCAGAGTTAAAGGCCT | pEAQ_NtCYP96T1 |
| Nar_162 | TCTGCCCAAATTCGCGAC | GATCACTTCGCACGCCGTAGCCAGCGTC | pEAQ_NtCYP96T1 |
|  | GACGCTGGCTACGGCGTGCGAAGTGATC | GAAACCAGAGTTAAAGGCCT | pEAQ_NtCYP96T1 |
| Nar_163 | TCTGCCCAAATTCGCGAC | GAAAGACACGGCATCGTAGC | pEAQ_NtCYP96T1 |
|  | ATAGCTACGATGCCGTGTCTT | GAAACCAGAGTTAAAGGCCTCGAGTCACATGACTGATCTCTTTCTAA | Nar_160 |
| Nar_164 | TCTGCCCAAATTCGCGAC | CATGGTTATCGATTGCGTGTTTCTCT | pEAQ_NtCYP96T1 |
|  | CGCAATCGATAACCATGACTTGGT | GAAACCAGAGTTAAAGGCCT | pEAQ_NtCYP96T1 |
| Nar_165 | TCTGCCCAAATTCGCGAC | CATGGTTATCGATTGCGTGTTTCTCT | pEAQ_Nar_160 |
|  | CGCAATCGATAACCATGACTTGGT | GAAACCAGAGTTAAAGGCCT | pEAQ_Nar_160 |
| Nar_166 | TCTGCCCAAATTCGCGAC | GTTTCTCTGCACTAACAAGTACGTGAA | pEAQ_NtCYP96T1 |
|  | GTACTTGTTAGTGCAGAGAAACACGCA | GAAACCAGAGTTAAAGGCCT | pEAQ_NtCYP96T1 |

**Supplementary Table 2:** Oligos used for cloning fragments for site-directed mutagenesis experiments. See Supplementary Table 3 for details on nomenclature of the clones. The two fragments were assembled into a pEAQ-HT digested vector using Gibson assembly (see methods). Underlined nucleotides indicate overlaps with vector.

| **Sequence Name** | **Mutant** |
| --- | --- |
| Nar_160 | NtCYP96T1 A217V VAWK251TACE A308L |
| Nar_161 | NtCYP96T1 A217V |
| Nar_162 | NtCYP96T1 VAWK251TACE |
| Nar_163 | NtCYP96T1 A308L |
| Nar_164 | NtCYP96T1 L315I |
| Nar_165 | NtCYP96T1 A217V VAWK251TACE A308L L315I |
| Nar_166 | NtCYP96T1 |

**Supplementary Table 3:** Nomenclature used for mutant sequences. Nar_160 was synthesized and used as a template.

| **Oligo** | **Sequence** |
| --- | --- |
| NtCYP96T1-F | TCTGCCCAAATTCGCGACCGGTATGGCCACTTCTTCTTCAG |
| NtCYP96T1-R | GAAACCAGAGTTAAAGGCCTCGAGTCACATGACTGATCTCTTTCTAA |
| NtSDR2-F | TCTGCCCAAATTCGCGACCGGTATGGAGAAGAAAAGGGTTTGTG |
| NtSDR2-R | GAAACCAGAGTTAAAGGCCTCGAGTCAATAACCAGCCTCCTTGT |
| NtCYP71DW1-F | TCTGCCCAAATTCGCGACCGGTATGCCACTGAATATTGTTTTGAGT |
| NtCYP71DW1-R | GAAACCAGAGTTAAAGGCCTCGAGCTAATCCTGCAGATTAACAGCC |
| NtODD1-F | TCTGCCCAAATTCGCGACCGGTATGATTGCCCCCTCTCAA |
| NtODD1-R | GAAACCAGAGTTAAAGGCCTCGAGCTACTCAAGAAGTACCATATTGTTTACC |
| NtOMT1-F | TCTGCCCAAATTCGCGACCGGTATGGGATCGTGCAATATTAATGA |
| NtOMT1-R | GAAACCAGAGTTAAAGGCCTCGAGCTACTTGTAGAACTCCATAATCC |
| NtODD2-F | TCTGCCCAAATTCGCGACCGGTATGGGTTCCGATTTCAAATCAC |
| NtODD2-R | GAAACCAGAGTTAAAGGCCTCGAGTTAGTTGTCGACTACTAGATTATTTA |
| NtCYP96T6-F | TCTGCCCAAATTCGCGACCGGTATGGTCACTTCTTCTTCAGC |
| NtCYP96T6-R | GAAACCAGAGTTAAAGGCCTCGAGTCACATGACTGATCGCTTTATAA |
| NtAKR1-F | TCTGCCCAAATTCGCGACCGGTATGTTGAACAACATTCCTGAG |
| NtAKR1-R | GAAACCAGAGTTAAAGGCCTCGAGTCATTCACCTTCATTTTCCTC |
| NtNMT1-F | TCTGCCCAAATTCGCGACCGGTATGATGGACGAACGGACCA |
| NtNMT1-R | GAAACCAGAGTTAAAGGCCTCGAGTTACTTTGGTTTATGACATGCAATG |
| NtCYP96T5-R | TCTGCCCAAATTCGCGACCGGTATGGCCACTTCTTCTCCATG |
| NtCYP96T5-F | GAAACCAGAGTTAAAGGCCTCGAGTTAGACGACTGATCTCTTCTTAA |
| NtSDR1-R | TCTGCCCAAATTCGCGACCGGTATGGAGAGGGCTGAGTAC |
| NtSDR1-F | GAAACCAGAGTTAAAGGCCTCGAGTCAACCGTTTATGCCCCG |
| NtCYP96T7-F | TCTGCCCAAATTCGCGACCGGTATGGCCACTTCTTCTTCAG |
| NtCYP96T7-R | GAAACCAGAGTTAAAGGCCTCGAGTTATATGGCTGATCTCTTTGTAAC |
| NtCYP96T8-F | TCTGCCCAAATTCGCGACCGGTATGGCCACTTCTTCTGCAG |
| NtCYP96T8-R | GAAACCAGAGTTAAAGGCCTCGAGTCACATGACTGATCTCTTTCTAA |
| NtCYP96T1_qPCR-F | TGCTATGGCGAGGATGAAGG |
| NtCYP96T1_qPCR-R | ACATGTCCCTTCACCATCTG |
| Histone_qPCR-F | GTCTGCCCCAACAACTGGAGG |
| Histone_qPCR-R | GCTTCCTAATCAGTAGCTCG |
| Vc_ActinQpcr_F2 | CCATGTATGTGGCTATCCAGGC |
| Vc_ActinQpcr_R2 | TCGAGACGAAGGATCGCATG |
| Vc_GABAT1_qpcr_F | CTGCTAGTCTCTCGGGCCTTA |
| Vc_GABAT1_qpcr_R | AGTTTCTCCAAATTATCAGCTAACCTTGACG |

**Supplementary Table 4:** Oligos used for cloning candidate genes. See Supplementary Table 3 for details on nomenclature of the clones. Underlined nucleotides indicate overlaps with vector.

1.	Breiterová, K., Koutová, D., Maříková, J., Havelek, R., Kuneš, J., Majorošová, M., Opletal, L., Hošťálková, A., Jenčo, J., Řezáčová, M., et al. (2020). Amaryllidaceae Alkaloids of Different Structural Types from Narcissus L. cv. Professor Einstein and Their Cytotoxic Activity. Plants *9*, 137. 10.3390/plants9020137.

2.	Kilgore, M.B., Augustin, M.M., May, G.D., Crow, J.A., and Kutchan, T.M. (2016). CYP96T1 of Narcissus sp. aff. pseudonarcissus Catalyzes Formation of the Para-Para’ C-C Phenol Couple in the Amaryllidaceae Alkaloids. Frontiers in Plant Science *7*.

3.	Liscombe, D.K., Usera, A.R., and O’Connor, S.E. (2010). Homolog of tocopherol C methyltransferases catalyzes N methylation in anticancer alkaloid biosynthesis. Proceedings of the National Academy of Sciences *107*, 18793–18798. 10.1073/pnas.1009003107.

4.	Liscombe, D.K., Louie, G.V., and Noel, J.P. (2012). Architectures, mechanisms and molecular evolution of natural product methyltransferases. Nat. Prod. Rep. *29*, 1238–1250. 10.1039/C2NP20029E.

5.	Huang, R., O’Donnell, A.J., Barboline, J.J., and Barkman, T.J. (2016). Convergent evolution of caffeine in plants by co-option of exapted ancestral enzymes. Proc Natl Acad Sci USA *113*, 10613–10618. 10.1073/pnas.1602575113.

6.	Choi, K.-B., Morishige, T., Shitan, N., Yazaki, K., and Sato, F. (2002). Molecular Cloning and Characterization of CoclaurineN-Methyltransferase from Cultured Cells of Coptis japonica*. Journal of Biological Chemistry *277*, 830–835. 10.1074/jbc.M106405200.

7.	Junker, A., Fischer, J., Sichhart, Y., Brandt, W., and Draeger, B. (2013). Evolution of the key alkaloid enzyme putrescine N-methyltransferase from spermidine synthase. Frontiers in Plant Science *4*.

8.	Liscombe, D.K., Usera, A.R., and O’Connor, S.E. (2010). Homolog of tocopherol C methyltransferases catalyzes N methylation in anticancer alkaloid biosynthesis. Proceedings of the National Academy of Sciences *107*, 18793–18798. 10.1073/pnas.1009003107.

9.	Tian, Y., Zhang, C., and Guo, M. (2015). Comparative Analysis of Amaryllidaceae Alkaloids from Three Lycoris Species. Molecules *20*, 21854–21869. 10.3390/molecules201219806.

10.	Augustin, M.M., Ruzicka, D.R., Shukla, A.K., Augustin, J.M., Starks, C.M., O’Neil-Johnson, M., McKain, M.R., Evans, B.S., Barrett, M.D., Smithson, A., et al. (2015). Elucidating steroid alkaloid biosynthesis in Veratrum californicum: production of verazine in Sf9 cells. The Plant Journal *82*, 991–1003. 10.1111/tpj.12871.
